# Scaling Large Language Models for Next-Generation Single-Cell Analysis

**DOI:** 10.1101/2025.04.14.648850

**Authors:** Syed Asad Rizvi, Daniel Levine, Aakash Patel, Shiyang Zhang, Eric Wang, Curtis Jamison Perry, Ivan Vrkic, Nicole Mayerli Constante, Zirui Fu, Sizhuang He, David Zhang, Cerise Tang, Zhuoyang Lyu, Rayyan Darji, Chang Li, Emily Sun, David Jeong, Lawrence Zhao, Jennifer Kwan, David Braun, Brian Hafler, Hattie Chung, Rahul M. Dhodapkar, Paul Jaeger, Bryan Perozzi, Jeffrey Ishizuka, Shekoofeh Azizi, David van Dijk

## Abstract

Single-cell RNA sequencing has transformed our understanding of cellular diversity, yet current single-cell foundation models (scFMs) remain limited in their scalability, flexibility across diverse tasks, and ability to natively integrate textual information. In this work, we build upon the Cell2Sentence (C2S) framework, which represents scRNA-seq profiles as textual “cell sentences,” to train Large Language Models (LLMs) on a corpus comprising over one billion tokens of transcriptomic data, biological text, and metadata. Scaling the model to 27 billion parameters yields consistent improvements in predictive and generative capabilities and supports advanced downstream tasks that require synthesis of information across multi-cellular contexts. Targeted fine-tuning with modern reinforcement learning techniques produces strong performance in perturbation response prediction, natural language interpretation, and complex biological reasoning. This predictive strength enabled a dual-context virtual screen that nominated the kinase inhibitor silmitasertib (CX-4945) as a candidate for context-selective upregulation of antigen presentation. Experimental assessment in human cell models unseen during training supported this prediction, demonstrating that C2S-Scale can effectively guide the discovery of context-conditioned biology. C2S-Scale unifies transcriptomic and textual data at unprecedented scales, surpassing both specialized single-cell models and general-purpose LLMs to provide a platform for next-generation single-cell analysis and the development of “virtual cells.”

## 1 Introduction

Single-cell RNA sequencing (scRNA-seq) has revolutionized our understanding of cellular heterogeneity by enabling the profiling of gene expression at single-cell resolution [1]. This technology has generated massive data atlases such as CELLxGENE [2] and the Human Cell Atlas [3], offering unparalleled opportunities for computational methods to extract biological insights from this data. Recent transcriptomic foundation models (FMs), such as scGPT [4], Geneformer [5], scFoundation [6], and scGenePT [7] have shown promise in modeling single-cell transcriptomic data at scale. Despite these advances, current models are often limited by custom architectures constrained to scRNA-seq data, hindering their scalability to larger model sizes, integration of different data modalities, and ability to perform diverse generative and predictive tasks. These limitations restrict the ability of expression-only foundation models to synthesize insights across datasets, modalities, and biological contexts, and highlight the opportunity for new approaches that can integrate diverse data types, including the rich contextual information contained in biological text and metadata.

Large Language Models (LLMs) [8, 9, 10] offer a promising solution to these challenges. Widely used in natural language processing (NLP), LLMs exhibit consistent performance improvements with scale across diverse downstream tasks [11, 12]. Their ability to process vast text corpora and generalize effectively to new applications makes them well-suited for addressing the limitations of current expression-only models. Cell2Sentence (C2S) [13, 14] provides a framework to leverage LLMs for biology by transforming high-dimensional single-cell data into a textual format. By converting scRNA-seq profiles into “cell sentences” – sequences of gene names ordered by expression level – C2S positions single-cell data within the LLM framework, providing better scalability and infrastructure advantages than specialized model architectures. This data transformation strategy simplifies model development and deployment, and enables easy integration of transcriptomic data with diverse modalities, including metadata, experimental conditions, and textual descriptions from biological publications.

Here, we introduce **C2S-Scale**, a new family of LLMs trained on a multimodal corpus of over 50 million cells and associated text. We show that scaling these models up to 27 billion parameters leads to consistent performance improvements across a range of predictive and generative tasks (Fig. 1). C2S-Scale’s flexible context allows it to analyze cellular interactions and diverse biological information in multi-cell contexts, enabling sophisticated applications from predicting perturbation responses to answering complex biological questions. To further enhance the biological accuracy of model outputs, we developed refinement techniques with reinforcement learning (GRPO) to align model predictions with key biological objectives. We also introduce a novel metric, single-cell Fréchet Inception Distance (scFID), for assessing generative performance.

**Figure 1:**
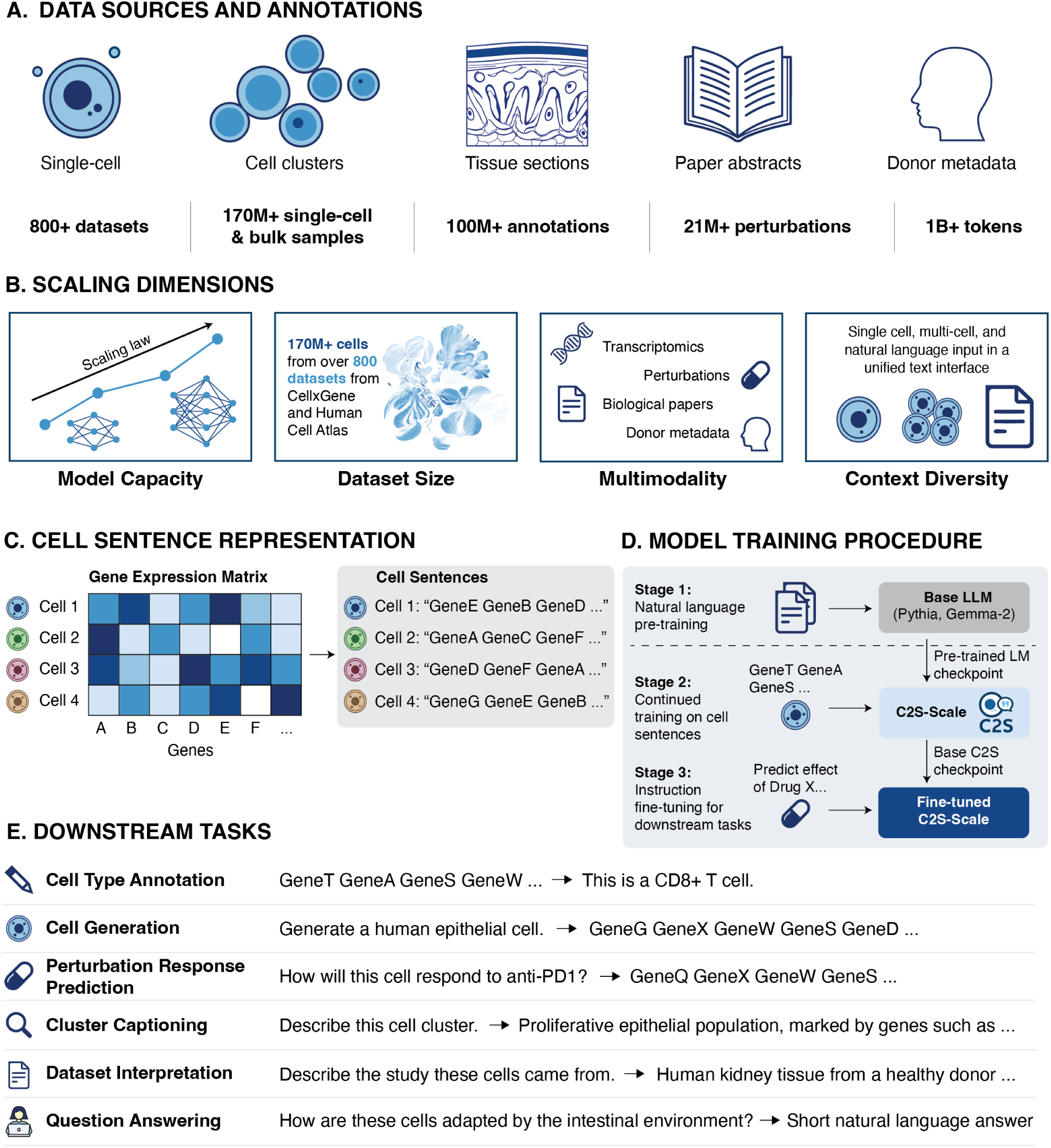
C2S-Scale unifies transcriptomics and natural language for scalable single-cell analysis. **A**, The C2S-Scale corpus integrates over 50 million single-cell transcriptomes curated from the CELLxGENE and Human Cell Atlas repositories, linking transcriptomic profiles with rich cell, tissue, and donor-level metadata as well as scientific abstracts and papers. **B**, C2S-Scale expands upon the Cell2Sentence framework by increasing model capacity up to 27 billion parameters, building a multi-task training corpus of over 1 trillion tokens, and expanding task diversity to include multimodal and multicellular inputs. **C**, Transcriptomic profiles are transformed into “cell sentences”, ordered sequences of gene names ranked by expression level. This enables the direct application of pretrained LLMs without requiring architectural modifications. **D**, C2S-Scale is initialized from pre-trained LLM checkpoints. An initial training phase on cell sentences is followed by instruction fine-tuning for a variety of downstream tasks. **E**, This unified training regime enables broad downstream capabilities, ranging from standard cell annotation to complex tasks like perturbation prediction and biological question answering.

To demonstrate the platform’s capacity for hypothesis generation, we programmed a dual-context virtual screen designed to prioritize potential context-conditional amplifiers of antigen presentation. The screen highlighted the kinase inhibitor silmitasertib (which has not been reported to enhance MHC-I expression) as a context-selective candidate, predicting a strong effect in the context of low levels of IFN exposure but minimal impact in the absence of IFN signaling. We experimentally evaluated this prediction in primary tumor fragments as well as independent human cell models unseen during training, where the results supported the model’s ability to identify biologically relevant, context-conditioned effects.

By releasing our models and resources, we provide a powerful, open-source platform for next-generation single-cell analysis.

## 2 Results

### 2.1 C2S-Scale: A unified framework for transcriptomics and natural language at scale

To bridge the gap between high-dimensional transcriptomic data and the reasoning capabilities of natural language systems, we developed **C2S-Scale**, a family of multimodal foundation models designed to interpret, predict, and generate single-cell biology at scale. We first assembled a massive, multimodal training corpus comprising over 50 million human and mouse transcriptomes curated from the CELLxGENE [2] and Human Cell Atlas [3] repositories. This corpus integrates diverse biological scales, linking single-cell profiles and multi-cell sample contexts with rich textual metadata and scientific abstracts (Fig. 1A).

Using this data, C2S-Scale expands upon the original Cell2Sentence framework [13, 14] by scaling along four major axes: model capacity, dataset size, multimodality, and context diversity (Fig. 1B). To leverage the extensive language capabilities of pretrained language models, we initialized C2S-Scale models from base pretrained Gemma-2 [15] and Pythia [16] checkpoints, and adapt them to the single-cell domain. This has the additional benefits of utilizing knowledge about biological concepts that the LLM picked up during pretraining, as well as utilizing the vast infrastructure developed around LLM training and inference. The unification in natural language is achieved by transforming gene expression profiles into “cell sentences”: ordered sequences of gene names ranked by expression level (Fig. 1C). The cell sentence transformation is reversible with minimal information loss (Supplementary Fig. 8). Unlike previous approaches that train custom architectures from scratch, C2S-Scale avoids modifying the underlying architecture of LLMs, instead focusing on a data transformation approach.

We perform an instruction finetuning phase of C2S-Scale models on multi-task single-cell analysis training samples (Fig. 1D), which we create by combining transcriptomic cell sentences with natural language instructions and textual annotations. This trains the models to perform a diverse set of objectives simultaneously through next token prediction (Table 1), enabling C2S-Scale to perform a broad spectrum of downstream tasks ranging from cell annotation to complex biological question answering and perturbation prediction (Fig. 1E). A key hypothesis of our work is that scaling model capacity unlocks emergent biological reasoning capabilities. To test this, we trained a spectrum of models ranging from 410 million to 27 billion parameters. This extensive scaling analysis reveals that larger models achieve higher performance and demonstrate superior multi-task capability. C2S-Scale effectively unifying single-cell analysis within a single, scalable computational framework in natural language.

**Table 1:** Pretraining tasks for C2S-Scale multi-task training. For multi-cell tasks, multiple cells are sampled from the same donor sample with the same tissue label.

| Task name | Type | Input information | Target output | Metric | Samples |
| --- | --- | --- | --- | --- | --- |
| Single cell language modeling | Single-cell | – | Single cell sentence | Overlap % | 28.8M |
| Cell type annotation | Single-cell | Single cell sentence | Cell type | BertScore | 12.0M |
| Conditional cell generation | Single-cell | Cell type of one cell | Single cell sentence | Overlap % | 24.7M |
| Multiple cell language modeling | Multi-cell | – | Multiple cell sentences | Overlap % | 24.7M |
| Tissue sample annotation | Multi-cell | Multiple cell sentences | Tissue label | BertScore | 10.2M |
| Sample cell type(s) annotation | Multi-cell | Multiple cell sentences | Cell types of multiple cells | BertScore | 10.2M |
| Conditional sample generation (tissue) | Multi-cell | Tissue annotation | Multiple cell sentences | Overlap % | 10.2M |
| Conditional sample generation (cell type) | Multi-cell | Cell types of multiple cells | Multiple cell sentences | Overlap % | 10.2M |
| Conditional sample generation (abstract) | Multi-cell | Paper abstract | Multiple cell sentences | Overlap % | 9.7M |
| Natural language interpretation | Multi-cell | Multiple cell sentences | Paper abstract | BertScore | 9.7M |
| Gene set enumeration | Gene set | Gene set name | List of genes in gene set | Overlap % | 27.6k |
| Gene set naming | Gene set | List of genes in gene set | Gene set name | BertScore | 27.6k |

### 2.2 C2S-Scale is a unified architecture for diverse single-cell analysis tasks

Current approaches to single-cell analysis often require piecing together different tools and architectures: specialized transcriptomic models for numerical tasks and separate LLMs for text-based interpretation. To demonstrate that C2S-Scale unifies these capabilities, we benchmarked our model against state-of-the-art expression-only foundation models (scGPT [4], Geneformer [5], scFoundation [6]), general-purpose LLMs (GPT-4o [17], Gemini [18]), and standard simpler models such as scVI [19] and linear baselines. We evaluated performance across three distinct task categories–predictive, embedding, and generative–which are summarized in Fig. 2A.

**Figure 2:**
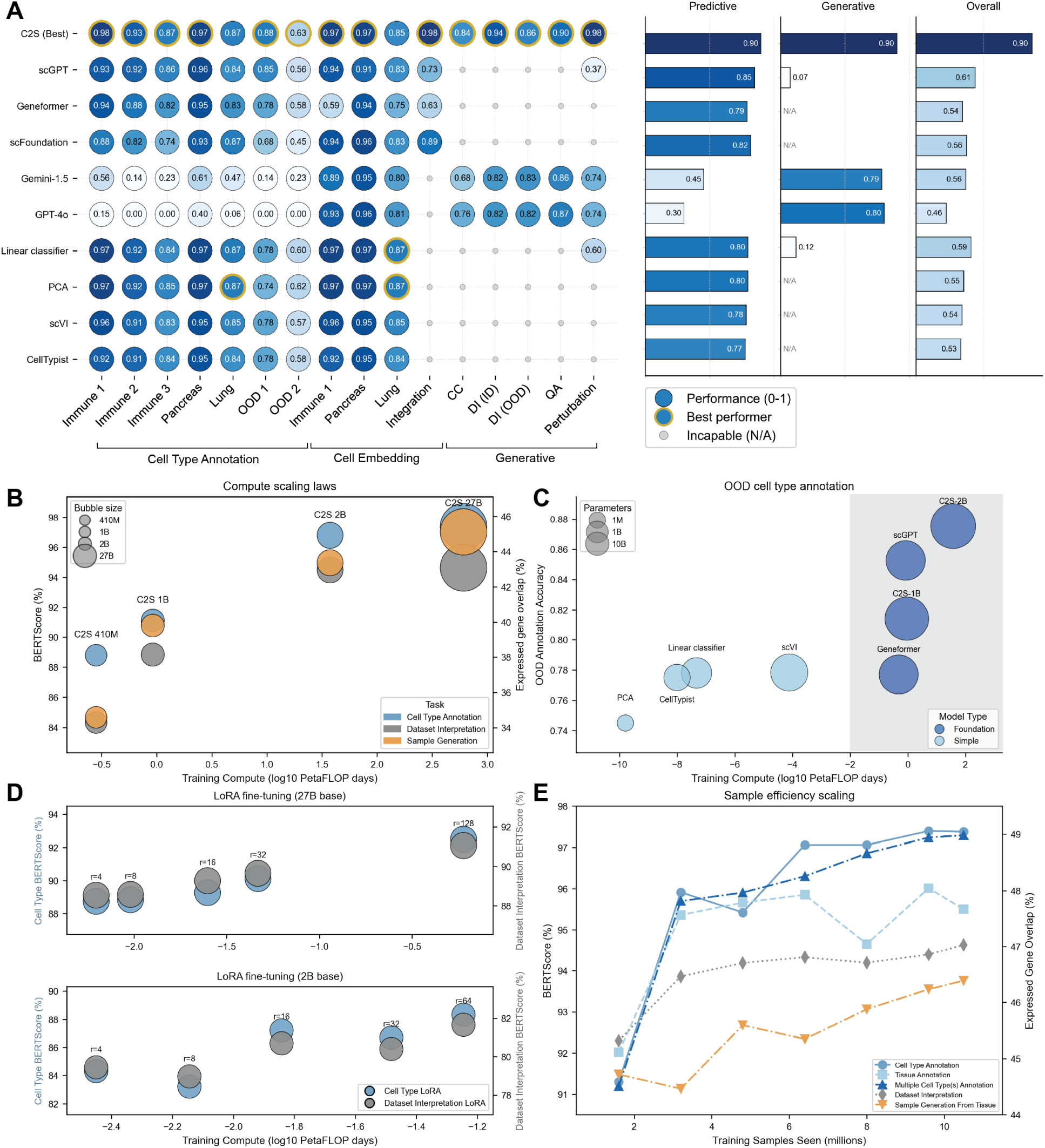
Caption continued on following page. C2S-Scale achieves state-of-the-art performance across diverse tasks and exhibits scaling behavior in performance. **A**, C2S-Scale outperforms expression-only foundation models (scGPT, Geneformer, scFoundation), general-purpose LLMs (Gemini, GPT-4), and simpler baseline models (PCA, linear classifier, etc) across a variety of tasks. Task names are shortened for brevity; see Supplementary Table 1 for the full length name. Cell type annotation and embedding tasks were evaluated using prediction accuracy against ground truth labels, natural language tasks were evaluated using semantic similarity (BERTScore), and perturbation response prediction was evaluated using scFID (reported as exp(−scFID) to normalize it to the same 0–1 range as the other tasks). The best model for each task is highlighted in gold. Aggregate scores for each task category were computed by averaging a model’s performance across all tasks in that category. C2S-Scale is the only model to demonstrate consistent, high-tier performance across all all tasks. **B**, Performance scaling as a function of training compute (FLOPs) for fully fine-tuned C2S-Scale models. As model capacity increases from 410M to 27B parameters, C2S-Scale shows consistent improvement in pretraining tasks such as cell type annotation, dataset interpretation, and sample generation. Cell type annotation and dataset interpretation are evaluated using BERTScore, while sample generation is evaluated as the percentage of generated genes in the true sample. **C**, Comparison of models for out-of-distribution (OOD) cell type annotation on immune cells. Foundation models, with larger parameter counts and training compute, show improved generalization in OOD settings compared to simpler baselines. **D**, Scaling trends persist in parameter-efficient fine-tuning regimes. Increasing the Low-Rank Adaptation (LoRA) rank yields consistent performance improvements, similar to the full-finetuned model. **E**, Sample efficiency scaling for the C2S-Scale-27B model. For a fixed model size, performance across tasks improves with the number of training tokens seen during pretraining.

On cell type annotation, C2S-Scale achieved results competitive with or superior to specialized expression-only models. Across several diverse immune cell datasets [20, 21, 22], pancreas [23], and lung [24] datasets, C2S-Scale annotates cells with *>* 85% accuracy, outperforming both foundation model baselines as well as simpler models. In a harder out-of-distribution cell type annotation task, where we train on samples from one dataset and evaluate on a different dataset, C2S retains high accuracy (88% on OOD scenario 1) while other models, especially non-foundation models, suffer drastically compared to an in-distribution setting. We discuss this setup in more detail in Section 2.4.

Embeddings taken from the final layer of C2S-Scale models also consistently show strong correspondence with cell annotations, outperforming embeddings from other methods on cell annotation based on embedding representations. Beyond annotation, we assessed the model’s ability to learn biologically meaningful representations through a multimodal integration task. Here, C2S-Scale was tasked with matching single-cell profiles to corresponding pseudo-bulk RNA-seq samples; notably, it accurately aligned these modalities despite having no prior exposure to bulk data, suggesting that the cell sentence representation captures robust, transferable cellular states.

Finally, C2S-Scale demonstrates unique strength in natural language interpretation and generation, a category of tasks that expression-only models are inherently incapable of performing. We evaluated interpretive performance using BERTScore to quantify semantic similarity between model outputs and ground truth annotations for each dataset and task. C2S-Scale outperformed leading general-purpose LLMs, including GPT-4o and Gemini-1.5, on tasks such as Cluster Captioning (CC, generating descriptive labels for cell clusters), Dataset Interpretation (DI, summarizing abstracts from data), and complex biological Question Answering (QA). These tasks are discussed in more detail in Section 2.4.

Taken together, these results confirm that C2S-Scale is a uniquely versatile platform, as demonstrated by its overall aggregated score across tasks (Fig. 2A). **To our knowledge, C2S-Scale is the only model capable of spanning this entire range of single-cell analysis tasks within a single unified architecture.**

### 2.3 Scaling enhances the biological reasoning capabilities of C2S-Scale

A central principle of modern LLMs is that performance improves predictably with increased scale [11, 12]. We tested this hypothesis for single-cell analysis by training C2S-Scale models ranging from 410 million to 27 billion parameters. We observed consistent scaling laws: as total training compute (FLOPs) and model size increased, performance improved across different pretraining tasks including cell type annotation, dataset interpretation, and conditional sample generation (Fig. 2B).

To evaluate whether increased computational cost yields tangible benefits over simpler methods, we analyzed the performance of C2S-Scale against expression-only foundation models (scGPT [4], Geneformer [5]) and standard baselines (scVI [19], CellTypist [20], linear models) on an out-of-distribution immune cell annotation task. While foundation models incur a higher computational footprint during pretraining, they demonstrate improved generalization capabilities compared to simpler baselines (Fig. 2C). This indicates that the computational investment in large-scale pretraining improves performance on unseen biological contexts. FLOPs usage by different models is given in more detail in Supplementary Table 1 and Supplementary Figure 9.

We also investigated scaling behavior in parameter-efficient training regimes. Using Low-Rank Adaptation (LoRA) [25] on the 2B and 27B parameter models, we found that performance on cell type annotation and dataset interpretation tasks improved consistently as the LoRA rank - and therefore the trainable capacity - increased (Fig. 2D). Performance also scaled positively with the volume of data observed; for the C2S-Scale 27B model, performance across pretraining tasks improved with the number of training tokens seen (Fig. 2E). These results confirm that scaling both model capacity and the diversity of the multimodal training corpus is a reliable strategy for enhancing biological reasoning.

### 2.4 Interpreting single-cell data across biological scales using natural language

Natural language interpretation is an underexplored aspect of single-cell analysis, enabling researchers to bridge experimental scRNA-seq data with existing biological literature and providing a user-friendly tool for biologists to interact with and interpret their data. Existing LLM-based single-cell models such as GenePT [26] and scGenePT [7] offered limited integration of natural language and single-cell data, focusing primarily on using language embeddings in single-cell architectures and tasks. C2S-Scale bridges large-scale training on transcriptomic data with the natural language pretraining and generative capabilities of LLMs, enabling natural language interpretation of scRNA-seq data at multiple scales of biology, illustrated in Fig. 3A.

**Figure 3:**
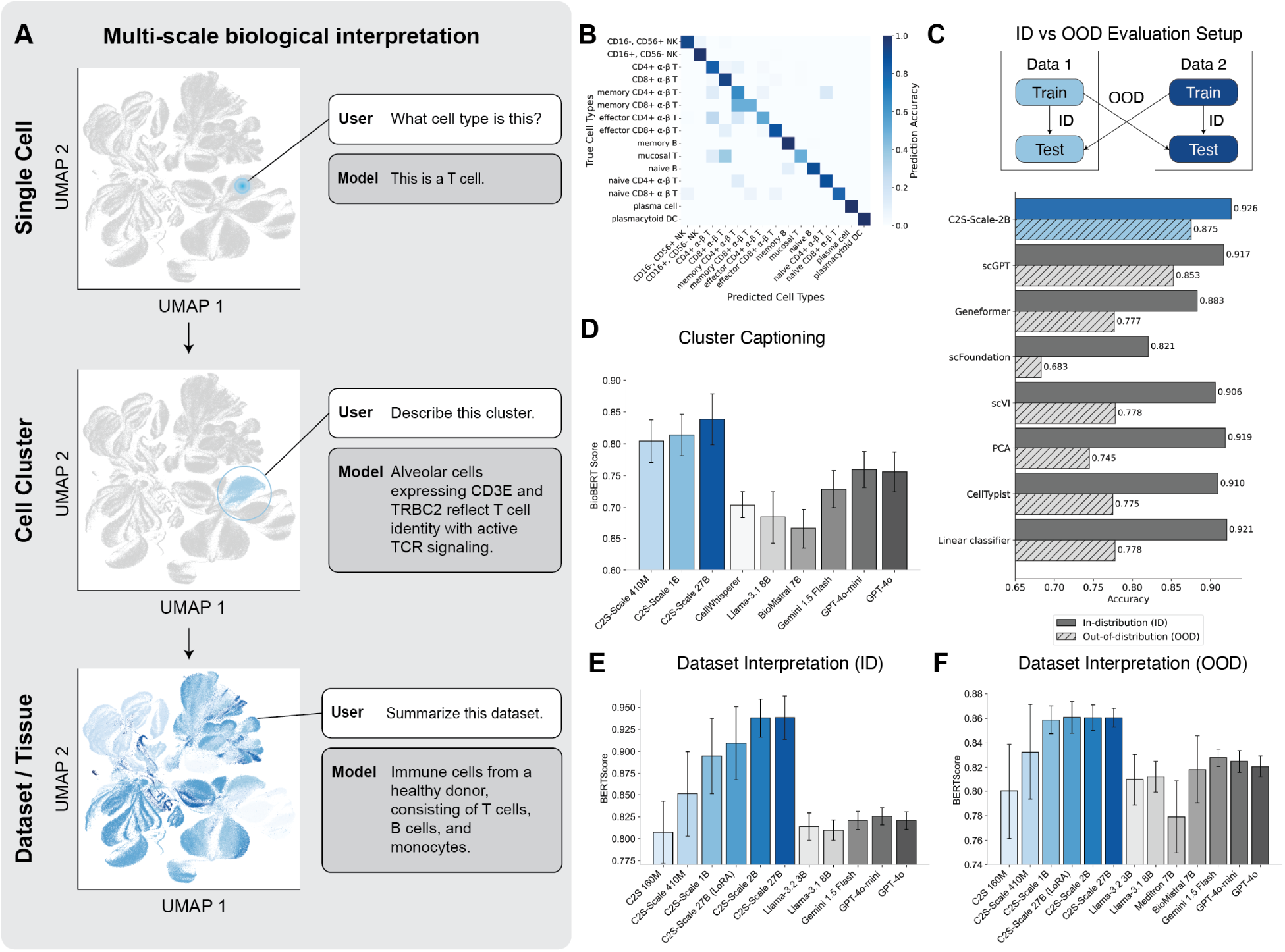
C2S-Scale enables natural language interpretation across biological hierarchies, from single cells to entire datasets. **A**, Overview of different scales of biological data interpretation, from single cells to cluster and dataset-level annotation. **B**, Single-cell annotation performance on a diverse immune cell atlas [21]. The correspondence between ground truth and predicted labels demonstrates accurate annotation at the single-cell level. **C**, Cell annotation performance of C2S-Scale versus foundation model and simple model baselines in an in-distribution versus out-of-distribution setting. C2S-Scale exhibits the smallest performance degradation when transferring from in-distribution (ID) to out-of-distribution (OOD) immune cell contexts. **D**, Cluster captioning performance on unseen data clusters. Models are tasked with generating descriptive biological captions given multi-cell context, with quality measured by BioBERTScore. **E-F**, Dataset-level summarization performance. Models generate natural language abstracts for entire scRNA-seq experiments, evaluated on held-out cells from seen studies (E) and completely unseen studies (F). Error bars represent standard deviation across test set samples.

We benchmark C2S-Scale on a series of natural language interpretation tasks at various scales of biology, evaluating its ability to reason about and generate meaningful descriptions about data. At the **individual cell level**, C2S-Scale is able to accurately annotate cell types in natural language given cell sentences as input. The model is first fine-tuned on a diverse immune cell dataset [21] to predict cell type labels in natural language. C2S-Scale is able to correctly classify almost all cell types on a held-out partition of the immune cell data (Fig. 3B), demonstrating C2S-Scale’s effectiveness at standard single-cell analyses. We also assessed the model’s robustness by evaluating performance on out-of-distribution (OOD) immune cells. Compared to both foundation model baselines and simpler linear models, C2S-Scale exhibited the smallest drop in performance when transferring to unseen biological contexts (Fig. 3C).

At the **cluster level**, we evaluate C2S on a Cluster Captioning task inspired by captioning tasks in computer vision, where the goal is to generate biologically meaningful descriptions for groups of cells from the same tissue and batch within a scRNA-seq dataset. To create training data for this task, we use GPT-4o to generate natural language captions for cell clusters derived from annotated datasets (Methods Section 4.7). C2S-Scale is fine-tuned to predict these captions given multiple input cell sentences from each cluster and is evaluated on held-out clusters not seen during training. Performance is measured using BioBERTScore [27], which quantifies semantic similarity between generated and ground-truth captions. As shown in Fig. 3D, C2S-Scale outperforms both baseline LLMs and CellWhisperer, another transcriptomic “captioning” model, on this task. Whereas CellWhisperer requires averaging the embeddings of the input cells, C2S is able to natively support multicellular input without aggregation, improving its ability to interpret and summarize nuanced expression patterns at the cluster level.

At the **dataset level**, we further evaluate interpretive ability through a Dataset Interpretation task, where the model receives multiple cell sentences from a scRNA-seq dataset and is tasked with generating a high-level summary in the style of a biological abstract. These summaries are expected to describe key features of the dataset, including dominant cell types, tissues, disease states, or perturbations (example provided in Fig. 13). Fig. 3E shows that C2S-Scale achieves the highest BERTScore among all evaluated models, including Llama [28, 29, 30], Meditron [31], BioMistral [32], Gemini [18], and GPT-4o [17]. Notably, C2S-Scale generalizes well to entirely unseen datasets, producing summaries that remain relevant and informative (Fig. 3F), highlighting its robust natural language understanding of scRNA-seq data.

Overall, C2S-Scale enables natural language interpretation at multiple scales, spanning single cells, clusters, and datasets. Its ability to integrate textual and biological data unlocks new opportunities for biologists to explore, annotate, and generate insights from scRNA-seq data in natural language.

### 2.5 C2S-Scale Learns Spatial Reasoning from Multi-cell Context and Interaction Data

Understanding spatial organization in tissues is fundamental to uncovering the mechanisms that govern cellular interactions, particularly in how they drive disease progression and tissue homeostasis [33, 34, 35]. Cellular niches, defined by their specific cell types, signaling molecules, and extracellular matrix components, play a crucial role in regulating these processes. Accurately predicting spatial relationships among cells from transcriptomic data alone is challenging, as traditional approaches often rely on explicitly structured spatial models or predefined interaction networks [36, 37, 38].

Although C2S-Scale was not explicitly designed for spatial reasoning, its ability to incorporate multi-cellular context provides a natural mechanism for modeling spatial organization. We hypothesize that by sampling and encoding cells from shared neighborhoods, C2S-Scale can infer spatial relationships without requiring architectural modifications. To test this, we evaluate the model’s performance in predicting spatial neighborhoods using a human liver spatial RNA-seq dataset [39]. Additionally, we simultaneously train C2S-Scale on related tasks aimed at improving its spatial understanding: niche label prediction, neighbor cell generation, and determining whether multiple cells belong to the same niche (Fig. 4A). By training on these complementary tasks, C2S-Scale learns robust representations of spatial organization, significantly outperforming Nicheformer [40], scGPT, GPT-4o, and a simple linear baseline across tasks (neighborhood prediction shown in Fig. 4B; additional tasks in Supplementary Fig. 17A-C). C2S excels in tasks that require understanding multicellular inputs, like neighborhood prediction. We find that Nicheformer succeeds at niche label prediction tasks, but struggles to synthesize multicellular information for more the more complex neighborhood prediction task..

**Figure 4:**
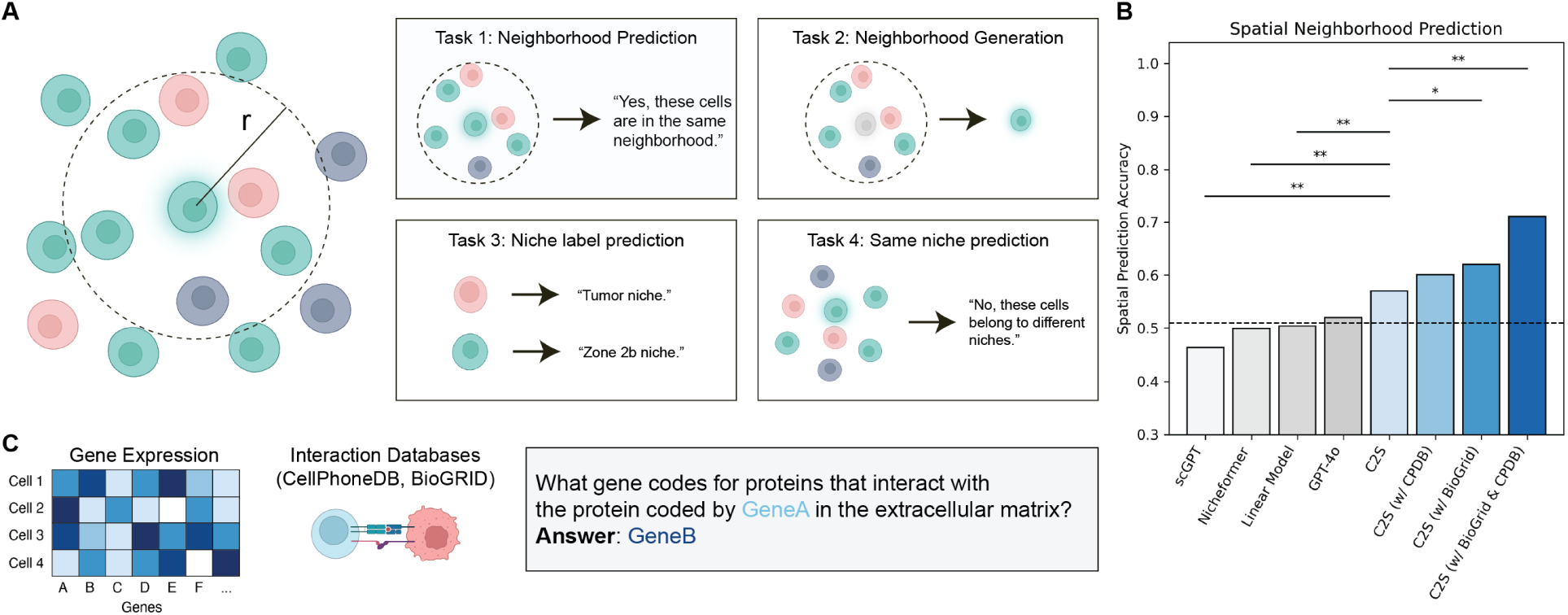
C2S-Scale can interpret multi-cellular spatial context and predict niche neighborhoods. **A,** We fine-tune C2S-Scale on a variety of single and multi-cellular spatial tasks designed to enable C2S-Scale to perform spatial reasoning, including identifying whether cells belong to the same neighborhood, generating spatial neighbors, and predicting the niche label of a cell or group of cells. A “neighborhood” is defined to be cells within a fixed radius from a central cell. **B,** C2S outperforms Nicheformer, scGPT, and other models in spatial neighborhood prediction accuracy (* = P < 0.05, ** = P < 0.01; McNemar’s test). Dashed line represents random baseline accuracy (0.51). **C,** We use publicly available gene interaction databases including BioGRID and CellPhoneDB to construct natural language interaction prompts about gene interactions. Integrating gene interactions from BioGRID and CellPhoneDB individually improves performance, and their combination provides the greatest improvement (Panel B). These results highlight the multi-task transfer learning potential of C2S-Scale for spatially-aware biological modeling.

We further hypothesize that incorporating external biological knowledge – specifically, gene interaction networks – can enhance spatial reasoning. Receptor-ligand and other protein-protein interactions are central to cell-cell communication, yet many scFMs are unable to integrate this information. Instead of imposing predefined rules, we simply expose C2S-Scale to receptor-ligand interactions from CellPhoneDB [41] and protein interaction data from BioGRID [42], formatted as natural language prompts (Fig. 4C). This approach allows the model to implicitly integrate prior knowledge while maintaining flexibility in how it applies this information.

Fine-tuning with gene interaction data further improves C2S-Scale’s ability to predict spatial relationships, reinforcing the hypothesis that external molecular context enhances spatial reasoning (Fig. 4B, Supplementary Fig. 17A-C). Notably, adding either CellPhoneDB or BioGRID data individually improves performance, demonstrating that both receptor-ligand and protein-protein interaction knowledge contribute to spatial reasoning (Fig. 4C). Moreover, combining both datasets results in the greatest improvement, suggesting that integrating diverse biological interaction sources allows LLMs to develop a richer understanding of multi-cellular organization and interactions. This improvement persists over different training runs of the C2S models with different random seeds (Supplementary Fig. 17D).

A key advantage of C2S-Scale is its ability to integrate diverse data sources without requiring explicitly structured incorporation of external knowledge. Unlike traditional methods that rely on predefined pathways or manually curated interaction models, C2S-Scale implicitly learns to incorporate relevant information during training. This highlights a fundamental strength of C2S: rather than designing bespoke architectures for specific tasks, we can provide relevant data, and the model autonomously determines how to utilize it. This capability extends beyond spatial reasoning and suggests broad applicability for integrating multimodal biological data.

### 2.6 Single-Cell Question Answering (QA) through Reinforcement Learning

QA tasks form a core part of NLP, providing a standard test to measure a model’s ability to understand information and apply reasoning [43, 44, 45, 46]. In biomedical research, QA tasks are particularly valuable for assessing advanced reasoning in domain-specific contexts, as evidenced by the development of numerous specialized QA datasets for medical [47, 48] and biological [49] applications. Building on this foundation, we introduce a single-cell Question Answering (scQA) task to assess the ability of foundation models to reason about and interpret single-cell transcriptomic data.

The scQA dataset consists of two thousand question-answer pairs, each containing: (i) an associated biological context, (ii) relevant scRNA-seq data sampled from clusters or cell type annotations, (iii) a main question, and (iv) a final answer. Additionally, each answer is annotated with keywords to help evaluate response quality. To construct the dataset, we sample cells from scRNA-seq datasets, provide the sampled data along with associated biological manuscripts to GPT-4.5 [17], and prompt it to generate meaningful questions (Fig. 5A).

**Figure 5:**
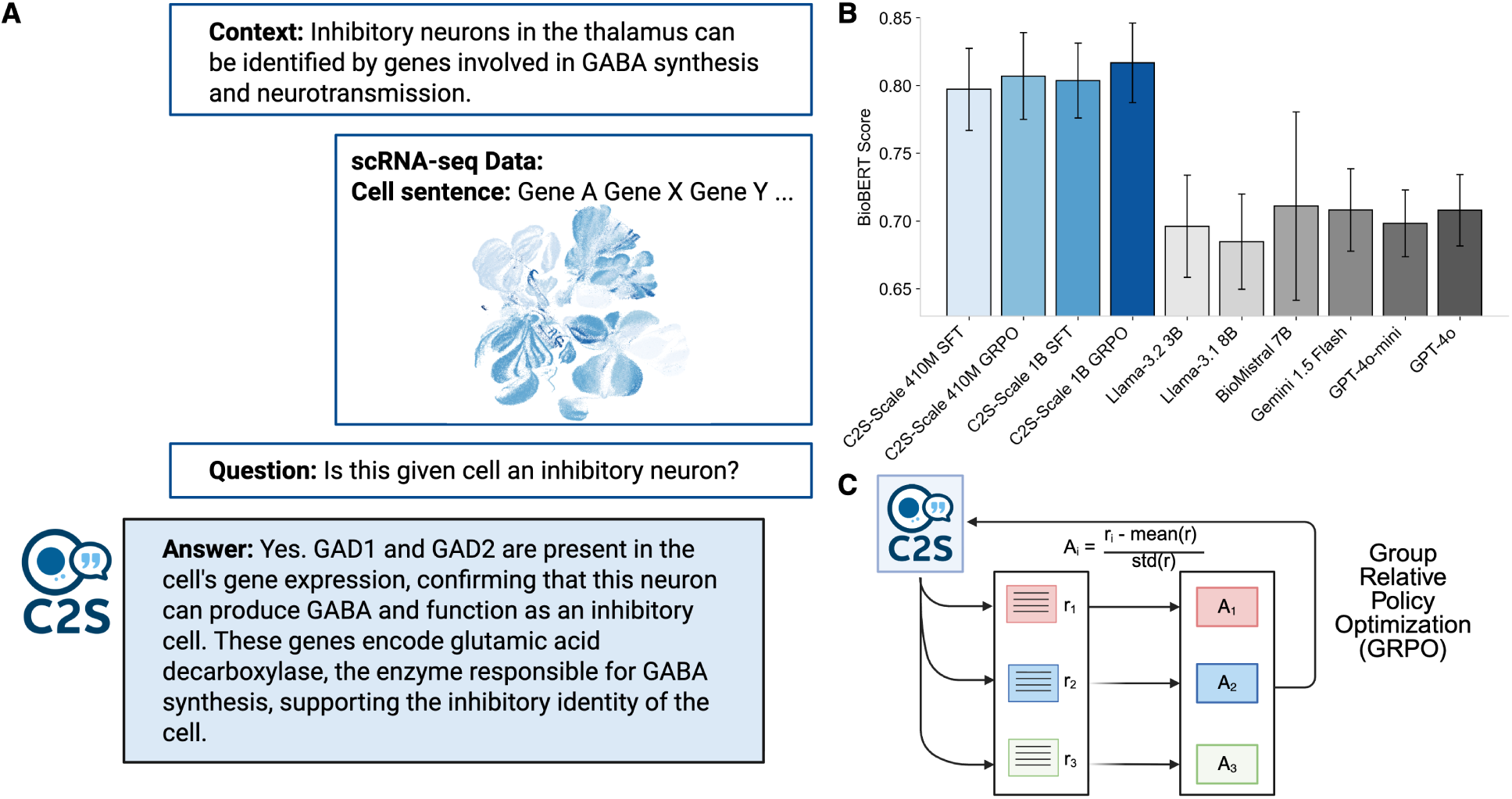
C2S-Scale demonstrates superior single-cell question answering performance compared to state-of-the-art (SOTA) LLMs. **A,** Example QA scenario based on scRNA-seq data. **B,** Empirical comparison of C2S-Scale and SOTA LLMs on single-cell QA tasks, highlighting C2S-Scale’s advantage in domain-specific reasoning. Error bars represent standard deviation across test set QA samples. **C,** Overview of the GRPO framework, which further refines model performance by training on preference data.

After supervised fine-tuning (SFT), C2S-Scale surpasses the performance of state-of-the-art LLMs on scQA (Fig. 5B), demonstrating the advantages of specialized training on transcriptomic data paired with natural language. To further improve C2S-Scale’s question answering capabilities, we employ Reinforcement Learning (RL) [50] through Group Relative Policy Optimization (GRPO) to further optimize the model to generate preferred responses to questions (Fig. 5C). By using BioBERTScore as the reward function, we guide C2S-Scale toward producing higher-quality answers aligned with biological insights. Following GRPO training, C2S-Scale significantly outperforms the SFT baseline on the scQA dataset, highlighting the potential of RL techniques to optimize LLMs for specialized single-cell applications.

### 2.7 Perturbation Response Prediction

Single-cell foundation models offer remarkable opportunities for conducting large-scale virtual perturbation experiments that would otherwise be infeasible or prohibitively expensive in a laboratory setting. Here, we demonstrate C2S-Scale’s generalization capabilities across unseen perturbations and cellular contexts, along with its broad applicability for modeling perturbation responses (Fig. 6A).

**Figure 6:**
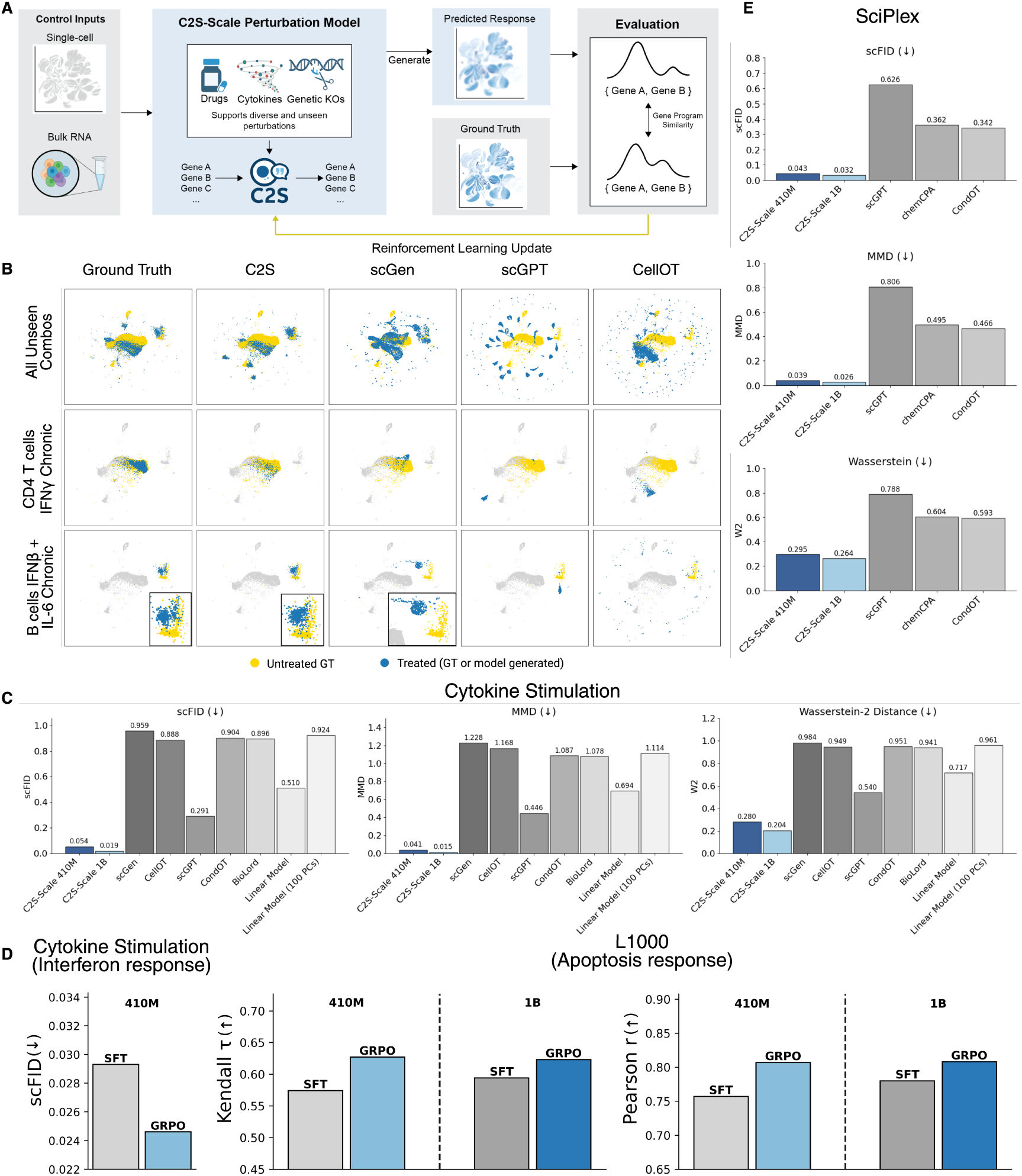
C2S-Scale models outperform existing methods in predicting cellular responses to unseen perturbations. **A,** Overview of the C2S-Scale perturbation prediction framework, which supports diverse perturbation types including drugs, cytokines, and genetic knockouts, utilizing Group Relative Policy Optimization (GRPO) to prioritize the accurate reconstruction of key gene programs such as apoptosis or interferon response. **B,** UMAPs comparing predicted vs. ground-truth responses for unseen perturbations across four models. Rows show: (1) all combinatorial perturbations, (2) CD4 T cells under IFN-*γ*, (3) B cells under the held-out IFN-*β* + IL-6 stimulation. C2S-Scale aligns closely with ground truth in all cases. **C,** Benchmark metrics show C2S-Scale outperforms scGen, CellOT, scGPT, CondOT, BioLord, and linear models across all evaluation criteria. **D,** GRPO improves over SFT on L1000 (apoptosis response) and cytokine stimulation (interferon response) tasks, with gains in Kendall’s *τ*, Pearson’s *r*, and scFID. **E,** Benchmarking on the SciPlex3 dataset for unseen small molecule perturbations. C2S-Scale outperforms ChemCPA, scGPT, and CondOT across scFID, MMD, and Wasserstein metrics, demonstrating the model’s ability to leverage pathway information for out-of-distribution prediction.

Training proceeds in two stages: supervised fine-tuning (SFT) to predict gene-expression profiles of untreated cells—including L1000 cell lines—under specified perturbation conditions, followed by online reinforcement learning with GRPO [51] that optimizes biologically relevant objectives. We designed the reward function to prioritize the accurate prediction of key gene programs of interest. This includes apoptosis for L1000 [52] and interferon response for Dong et al. [53]. Concretely, the reward is computed over these targeted gene subsets (Fig. 6A), which focuses optimization while preserving full-profile generation and improves out-of-distribution generalization.

We introduce a new metric, scFID (Supplementary Fig. 19A), an adaptation of the FID metric [54] widely used in computer vision to evaluate the realism of generated images. scFID adapts the FID metric by replacing the Inception-v3 model with a single-cell foundation model to embed transcriptomic data, enabling evaluation of generated cells in a representation space aligned with biological structure and functional gene programs. By assessing differences in this embedding space rather than at the level of individual genes, scFID captures higher-order variation across cell states, yielding stable model rankings (Supplementary Fig. 19B-G) and aligning with distributional similarities evident in cell-state embeddings, while complementing expression-level metrics such as Kendall’s *τ* and Pearson’s *r*.

C2S-Scale outperforms existing methods on the Dong et al. dataset, accurately predicting responses to unseen cytokine perturbations on entire gene expression profiles. It generalizes to novel combinations of cell type, cytokine, and exposure duration, highlighting its ability to transfer to completely new contexts not seen during training (Fig. 6B). Compared to baselines, C2S-Scale performs best on fully unseen combinatorial perturbations, capturing nonlinear synergistic effects. Quantitative results (Fig. 6C) show superior MMD, Wasserstein, and scFID scores relative to competing models. In particular, both C2S-Scale and scGPT [4] outperform the linear models, showing the superiority of foundation models in predicting responses to unseen perturbations. GRPO further reduces scFID on interferon-related genes by 16%, thereby improving biological fidelity in immune pathways (Fig. 6D, left).

The L1000 results further underscore C2S-Scale’s versatility in modeling perturbation responses across single-cell and bulk transcriptomic data. We evaluate performance on apoptosis-related genes, focusing on generalization to unseen compound treatments. Applying GRPO yields consistent gains (Fig. 6D, right), improving Kendall’s *τ* by 9.2% for the 410M model and 4.9% for the 1B model, and Pearson’s *r* by 6.6% for the 410M model and 3.6% for the 1B model. Rewards are defined on phenotype-linked gene programs (e.g., apoptosis in L1000 [52] and interferon response [53]), which yields context-aware scores well suited for virtual screening and candidate prioritization, while preserving full-profile prediction and enhancing out-of-distribution generalization.

Finally, we evaluated performance on the SciPlex3 dataset [55], which presents a more difficult generalization challenge than the cytokine task. In this setting, the test set comprises cells treated with small molecule perturbations that were completely unseen during training, necessitating that models leverage molecular properties to generalize. Despite this increased difficulty, C2S-Scale significantly outperforms competing baselines, including ChemCPA [56], scGPT, and CondOT (Fig. 6E). We observe superior performance across scFID and MMD metrics (with Wasserstein distance results detailed in the supplement), demonstrating the model’s capacity to effectively translate molecular and pathway information into accurate transcriptomic predictions for novel drug treatments.

### 2.8 Virtual screening with C2S-Scale nominates a novel candidate for amplifying antigen presentation

A compelling application of single-cell perturbation models lies in their potential to accelerate drug repurposing through high-throughput virtual screening. By simulating responses to a vast array of compounds, these models enable rapid prioritization of therapeutic candidates *in silico*, bypassing the resource constraints of physical screening. This capability is particularly valuable for identifying agents that act conditionally within specific biological contexts, such as the tumor microenvironment, where standard culture conditions may fail to capture relevant therapeutic mechanisms.

To evaluate whether C2S-Scale can leverage this capability to prioritize context-dependent determinants of immune visibility, we applied a dual-context *in silico* screen (Fig. 7A). We focused specifically on identifying compounds that enhance MHC-I antigen presentation in an immune-context-positive state (defined by primary tumor profiles with high baseline type-I interferon signaling) while showing minimal activity in a neutral state (defined by cell line profiles with low baseline type-I interferon signaling).

**Figure 7:**
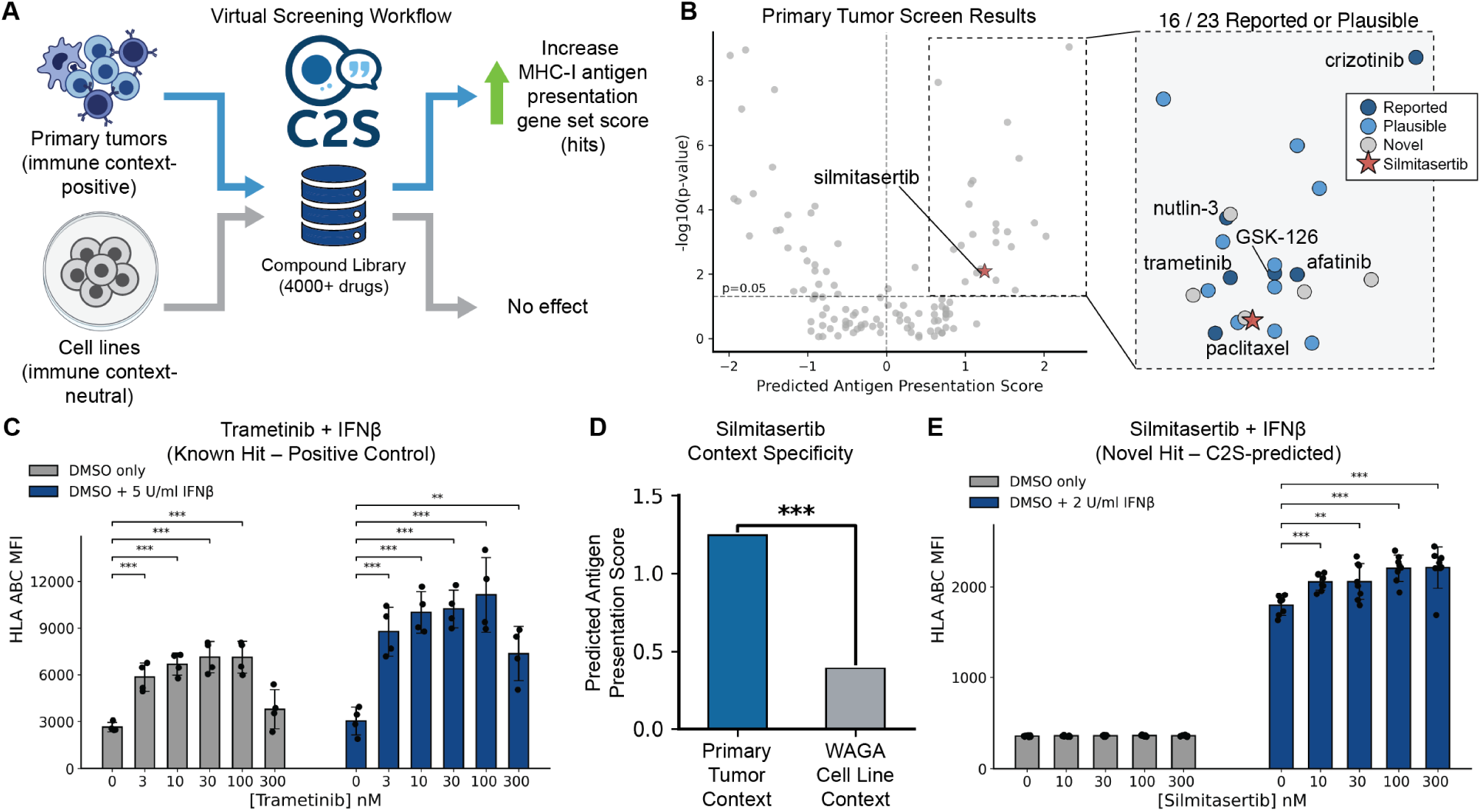
Virtual screening with C2S-Scale nominates a context-dependent modulator of antigen presentation. **A,** Schematic of the dual-context virtual screening workflow. The model predicts perturbation outcomes on MHC-I antigen presentation gene sets using initial states derived from primary tumor profiles (immune context-positive) and cell line profiles (immune context-neutral). **B,** Volcano plot displaying predicted drug effects in the primary tumor screen. Silmitasertib (red star) is identified as a high-scoring novel candidate. Inset details the top-ranking compounds, stratified by prior evidence: known MHC-I modulators (dark blue), plausible candidates with pathway links (light blue), and novel hits (grey). **C,** Experimental validation of the positive control, trametinib. Dose-response assessment of HLA-A,B,C surface levels (Mean Fluorescence Intensity; MFI) in WAGA cells treated with indicated concentrations of trametinib in the absence (gray) or presence (blue) of IFN-*β* (2 U/ml). **D,** Predicted context specificity of silmitasertib, showing a significantly higher antigen presentation score in the primary tumor context compared to the cell line context. **E,** Experimental validation of the novel hit, silmitasertib. Dose-response assessment of MHC-I MFI (normalized to vehicle control) in WAGA cells treated with silmitasertib without (gray) or with (blue) low-dose IFN-*β* (2 U/ml). Error bars represent mean ± s.d. Significance calculated via two-way t-test with Benjamini-Hochberg correction for multiple comparisons; ∗*P <* 0.05, ∗ ∗ *P <* 0.01, ∗ ∗ ∗*P <* 0.001, ∗ ∗ ∗ ∗ *P <* 0.0001.

The screening results provided initial validation of the model’s predictive power by recovering established biology. The screen prioritized several compounds reported in the literature to amplify antigen presentation, including the MEK inhibitor trametinib, which appeared as top hits in the primary tumor context (Fig. 7B, dark blue markers). Analysis of the top-ranked candidates revealed that 16 of the top 23 hits were either reported modulators or mechanistically plausible (Fig. 7B, inset). To test the model’s ability to discover novel candidates, we examined high-scoring compounds that lacked prior literature evidence for this specific indication. We selected silmitasertib (CX-4945), a CK2 inhibitor, for experimental validation (Fig. 7B, red star). A detailed list of the top hits and their literature status is provided in Supplementary Table 3. To validate these screening results in the relevant biological setting, we first tested a panel of top-scoring candidates in an immune context-positive tumor fragment model (Supplementary Fig. 18A). We observed statistically significant upregulation of MHC-I surface levels for 5 of 6 predicted hits, including both established modulators and the novel candidate silmitasertib.

To further investigate the predictions generated by the screen, we utilized cell line models to test the impact of specific environmental factors. We hypothesized that the “immune-context-positive” state relied on inflammatory features, specifically type I interferon signaling, which is a hallmark of the tumor microenvironment often absent in standard cell culture [57, 58]. To benchmark the model’s performance against a known positive control, we first tested trametinib in A375 melanoma cells, as it has previously been studied in BRAF-mutant contexts [59, 60]. Trametinib treatment yielded a robust, dose-dependent increase in HLA-A,B,C surface levels both in the absence and presence of exogenous interferon-*β* (Fig. 7C). This confirms the model’s ability to identify potent modulators that maintain efficacy across varying immune contexts.

To characterize the “immune-context-neutral” state, we computationally screened three cell lines not included in the C2S-Scale training data: WAGA (Merkel cell), MCC2 (Merkel cell), and UTSCC45 (head and neck squamous cell). Analysis of baseline gene signatures confirmed that all three lines exhibit low interferon response scores compared to the primary tumor context (Supplementary Fig. 18B). The model predicted no significant MHC-I upregulation for silmitasertib across the three cell lines (Fig. 7D, Fig. 18B). We selected the WAGA line for experimental validation to test whether reconstituting this specific environmental factor could rescue the therapeutic effect. As predicted by C2S-Scale, silmitasertib treatment alone had no effect on MHC-I surface levels. However, the combination of silmitasertib with low-dose interferon-*β* (mimicking the immune-context-positive state) resulted in a reproducible, dose-dependent increase in MHC-I mean fluorescence intensity (MFI) (Fig. 7E). This amplification effect was preserved in MDK-knockout WAGA cells (Supplementary Fig. 18C) and was not restricted to a specific interferon subtype, as silmitasertib also enhanced MHC-I expression in the presence of low-dose interferon-*γ* (Supplementary Fig. 18D).

These results illustrate the potential of C2S-Scale to effectively sift through massive perturbation libraries to identify promising novel therapeutic candidates for experimental validation. While the precise biological mechanism by which silmitasertib amplifies the interferon response requires further characterization, the alignment between the model’s context-specific predictions and the experimental data exhibits the utility of C2S-Scale for high-throughput virtual screening.

## 3 Discussion

Although artificial intelligence approaches including neural network models have achieved significant breakthroughs in protein structure and the prediction of molecular interactions, less progress in modeling multi-cellular tissues, pathologic states, and context-specific biology has been made. Principal challenges in this space include the underlying diversity, complexity, and pleiotropy of biological systems, which compounds across hierarchical organization from genes to transcriptional programs, and cells to tissues to organisms. Indeed, the semantic complexity and contextuality of biological systems seems unrivaled–outside of language itself. Our work introduces C2S-Scale, a family of LLMs for single-cell analysis that leverages the benefits of state-of-the-art LLMs out of the box. By converting transcriptomic profiles into “cell sentences,” C2S-Scale avoids the need for bespoke model architectures while readily integrating contextual information from annotations, metadata, and biological texts. This data engineering paradigm yields a flexible system capable of predictive and generative single-cell tasks, and our results demonstrate that scaling C2S-Scale up to 27 billion parameters systematically boosts performance, mirroring similar scaling phenomena observed in the broader field of NLP.

Moreover, we show that C2S-Scale bridges the gap between raw transcriptomic information and natural language-based interpretation by supporting tasks at multiple scales, ranging from cell type annotation to entire dataset-level summarization. We propose new evaluation datasets for these interpretation tasks and demonstrate that LLMs trained in the C2S-Scale framework provide meaningful captions and summarizations of single-cell data, even in cases where the dataset is completely new to the model. By aligning expression data with rich textual metadata and biological domain knowledge, our approach highlights the potential of language-based modeling to offer biologically informed explanations and generate insights unavailable to purely expression-only systems.

Context-specific decoding is a core task for both LLMs and biological systems alike. To test the ability of C2S-Scale to derive context-specific biological meaning, we conducted a conditional virtual screen, identifying an IFN-specific regulator of antigen presentation. We validated the effectiveness of silmitasertib in neuroendocrine Merkel cell models and primary tumor models in which the downregulation of antigen presentation machinery is a well-established mechanism of resistance to immunotherapies. This success provides a blueprint for future screens targeting other complex biological contexts.

We anticipate that higher-capacity models and more diverse training corpora will unlock advanced capabilities, such as the integration of epigenomic, proteomic, and clinical data into a single multimodal model. In parallel, increasing transparency and explainability in LLM decision making will be essential for building trust and accelerating adoption of these tools in single-cell research. Reinforcement Learning and other innovations in LLM alignment will provide a path forward for aligning LLMs to preferred responses in the context of biological tasks. By directly linking natural language and transcriptomic data, C2S sets the stage for transformative innovations in biological discovery and personalized medicine.

## 4 Methods

The following section details the data collection, processing, and formatting for multi-task samples, as well as the model architecture for Large Language Models.

### 4.1 Data Collection

To construct the C2S-Scale pretraining corpus, we assembled over 50 million single-cell transcriptomic profiles from human and mouse tissues gathered from 825 scRNA-seq datasets (dataset links provided in Supplementary Table 2). Datasets were obtained from established public repositories, including the CELLxGENE [2] and Human Cell Atlas [3] data portals, and span a wide range of tissues, disease states, and experimental conditions. Each dataset was accompanied by author-provided metadata, such as cell type and tissue annotations, donor information, developmental stage, and associated study identifiers. A comprehensive summary of cell types, tissues, organisms, assays, diseases, sexes, and ages for the final corpus are provided in Supplementary Table 4. Where available, supplementary textual resources, including paper abstracts and study descriptions, were also retained.

Following established conventions [61], we preprocess raw scRNA-seq data by removing genes that are expressed in fewer than three cells and cells that express fewer than 100 genes, followed by normalizing counts to 1 × 10^4^ and log-transforming the data. Notably, we do not do any quality control for mitochondrial genes, as we found that proportions varied significantly across datasets and a single threshold was either too permissive or too restrictive across the corpus. For each dataset, the transcriptomic profiles were converted into cell sentences, and the accompanying annotations were preserved to construct natural language prompts. This resulted in a multimodal corpus linking expression profiles with textual descriptors of biological context.

Training and test sets for the pretraining corpus were created by splitting out 10% of cells from each scRNA-seq study and reserving them for the test set. This resulted in a held-out split of cells per dataset that reflects the tissue diversity seen within the training set. For harder out-of-distribution generalization scenarios, such as in dataset interpretation or cell annotation on unseen datasets, we download newer datasets that were released after the original corpus creation date, to prevent leakage in the training set and ensure a held-out dataset evaluation.

### 4.2 Cell Sentence Transformation

To adapt high-dimensional single-cell gene expression data into a format compatible with natural language processing, we converted expression profiles into textual representations termed “cell sentences.” For each cell, let *X* ∈ R*^D^* be the expression vector, where *X_k_*denotes the normalized expression value of gene *k* in that cell. The cell sentence for *X* is constructed by rank-ordering the genes within a cell by their expression levels and taking the *K* most highly expressed genes. If *S* is a list of indices from 1 to *D* sorted in descending order based on expression level in *X*, then

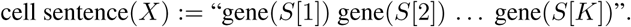

The gene names are in natural language, forming a sentence interpretable by language models (exemplified in Fig. 1C). In the case of ties in expression value, we break ties randomly to avoid introducing patterns in ordering between different genes with the same expression. In practice, we select *K* depending on the task the model is performing when constructing cell sentences and prompts in order to optimize memory and compute usage of our model. For predictive tasks such as cell type annotation on single cells, we find that *K* = 1000 often works well for cell annotation, achieving performance competitive or better than baseline models. For multi-cell context tasks such as cluster captioning or tissue annotation, we sample up to 20 cells for one input sample to C2S-Scale, and pick *K* such that the total number of tokens for the cells combined is close to the model’s context length (default is 8192 tokens), with each cell represented by *K* genes. We allow for slight variation in *K* for different samples containing the same number of cells, so that the model does not overfit to seeing a particular number of cells for certain numbers of cells. For generation tasks such as perturbation prediction, where we generate entire response cells, we set *K* to the length of the full cell sentence so that the model learns to generate the entire cell.

Under this framework, we do not extend or modify the vocabulary of the language model to include genes, instead opting to allow the LLM to tokenize genes and represent them using the existing token vocabulary it learned during natural language pretraining. This has two primary benefits: (i) by avoiding architectural modifications, the C2S framework is immediately applicable to any LLM architecture or innovation, and (ii) the LLM is able to recognize gene names and associate prior knowledge about that gene obtained during self-supervised pretraining on natural language data, which has been shown to be significant for large-scale pretrained LLMs [26]. We do not introduce any new special tokens to format cell sentences, and instead focus on prompt formatting to integrate cell sentences naturally as a part of LLM input.

The cell sentence transformation into textual sequences retains the underlying biological information by preserving the rank-order of gene expression. We find there is a strong linear relationship (in log space) between a gene’s rank in the cell sentence and the (normalized) expression level, validating the fidelity of this transformation. This relationship is shown in Supplementary Fig. 8 for two scRNA-seq datasets. A linear model fitted between rank and original expression can predict the original gene expression values given a gene’s rank with *R*^2^ = 0.85, demonstrating that minimal information is lost during conversion to cell sentences. This interchangeability allows us to utilize the strength of LLMs in natural language processing while retaining the ability to convert back to gene expression vectors for traditional single-cell analysis methods. The parameters of the linear model for each scRNA-seq dataset used during training are saved to enable reversible transformation from cell sentences back to expression values during inference.

### 4.3 Multi-Task Dataset Creation

C2S-Scale was designed to operate in natural language, enabling a broad range of tasks in single-cell analysis. These tasks include cell type and tissue annotation, multi-cell generation, and dataset-level interpretation. The complete list of pretraining tasks, is provided in Table 1, along with the number of training samples per task in the training corpus.

Prompts were constructed by combining the cell sentence representation of one or more cells with task-specific natural language instructions. For predictive tasks, the input prompt included a cell sentence and an instruction, and the output corresponded to the metadata label of interest. For example, in the cell type annotation task, the input consisted of the cell sentence and the instruction “Predict the cell type of this cell”, and the output was the corresponding cell type label. For generative tasks, this structure was inverted: metadata conditions were provided in the input prompt, and the model was trained to generate one or more cell sentences in response. Given that the maximum context length of C2S-Scale models was 8192 tokens, we adjusted cell sentence formatting per task to ensure that samples fit within the context window of each model: for single-cell tasks, we truncated the cell sentence if it exceeded 8000 tokens (around 2k genes), keeping the top *K* expressed genes, and for multi-cell tasks we ensured that the total tokens across *K* genes per each cell did not exceed the context length. Examples of these prompts for each task are available in Supplementary Figures 10, 11, 12, 13, 14, 15, 16.

Metadata included in natural language prompts encompassed cell type, tissue annotations, perturbation conditions, disease states, and text from associated studies or abstracts, thereby providing additional biological context. This framework enables C2S-Scale to interpret instructions, integrate biological knowledge, and generalize across diverse applications.

### 4.4 C2S-Scale architecture and pretraining

C2S-Scale employs large language models (LLMs) based on the Transformer architecture [8] to model cell sentences in natural language. The following sections go through the architectural components and training objectives of the transformer model architecture.

#### 4.4.1 Input representation

Input sequences are represented as high-dimensional embeddings suitable for processing by neural networks. Each word in a cell sentence corresponds to a gene name, which is first tokenized using the pretrained tokenizer associated with the backbone model. This approach avoids the introduction of new vocabulary and maintains compatibility with the LLM’s pretraining knowledge.

Tokenized gene names are mapped into vector representations through an embedding layer trained alongside the model. These embeddings capture semantic properties of genes informed both by their biological context and by the pretrained model’s prior knowledge. Positional encodings are added to preserve the rank order of genes within each cell sentence, allowing the model to learn dependencies across expression-ranked sequences.

#### 4.4.2 Attention mechanism

The central component of the Transformer is the self-attention mechanism [62, 8], which enables the model to compute pairwise relationships between tokens. For single-cell tasks, this allows the model to dynamically prioritize genes that are most informative for a given context, such as lineage-defining markers for classification or perturbation-responsive genes for prediction. The attention mechanism also extends naturally to metadata tokens (e.g. cell type, tissue, disease state), enabling the model to integrate gene expression with contextual information in a shared representation.

#### 4.4.3 Model architecture

C2S-Scale adopts a decoder-only Transformer design [17], chosen for its capacity to model sequential data and support generative tasks. The architecture consists of a stack of Transformer blocks, each containing a multi-head self-attention layer followed by a position-wise feedforward network. Residual connections and layer normalization are applied throughout to stabilize optimization and facilitate scaling to billions of parameters. This modular structure allows the model to capture long-range dependencies in gene expression data while remaining computationally efficient.

#### 4.4.4 Pretraining objective

The model is pretrained with a next-token prediction objective [63], in which the model learns to predict the next token in a sequence given all preceding tokens. Applied to cell sentences, this involves predicting the next gene in the rank-ordered expression list, optionally conditioned on metadata tokens. This autoregressive formulation encourages the model to capture the hierarchical organization of gene expression programs and to integrate biological context during generation.

In contrast to masked-token objectives such as those used in Geneformer [5], which predict randomly masked genes in non-linguistic sequences, the autoregressive objective aligns naturally with downstream generative applications. Training the model in this way conditions it to produce coherent, biologically meaningful outputs for tasks such as cell generation, dataset-level interpretation, and question answering.

#### 4.4.5 Training Setup

The main training phase of C2S-Scale models (Fig. 1D, Stage 2) was carried out on the C2S-Scale training corpus of more than 50 million single-cell transcriptomes with associated metadata and textual annotations, constructed as described in Section 4.1 and formatted into multi-task training samples as described in Section 4.3. C2S-Scale models are initialized from the base LLM weights pretrained on natural language data [16, 15], and undergo the main continued training phase on the C2S-Scale multi-task training corpus to adapt to the single-cell domain. Training was performed using the standard autoregressive next-token prediction objective implemented via established LLM training libraries, with loss taken over the entire sequence (prompt and response) to facilitate learning both the distribution of transcriptomic data represented by cell sentences as well as how to use cell sentences to perform different tasks.

Training infrastructure varied by model scale to accommodate computational requirements. The C2S-Scale 410M and 1B models were trained at Yale University using the Huggingface Transformers library (version 4.46.3) [64] and PyTorch (version 2.4.1) [65] on a High Performance Computing (HPC) cluster running Red Hat Enterprise Linux release 8.10. Each of these models was trained on 1-2 Nvidia H100 GPUs for several days (exact compute duration listed in Supplementary Table 1). The larger C2S-Scale 2B and 27B models were trained at Google using the JAX library. The C2S-Scale 2B model was trained on 288 TPU v4s for approximately 70 hours, while the C2S-Scale 27B model was trained on 272 TPU v5es for approximately 224 hours. Detailed statistics regarding FLOPs and compute usage are provided in Supplementary Table 1.

### 4.5 Scaling Evaluation

To evaluate scaling behavior, we analyzed the performance of C2S-Scale models immediately following the main training phase on the multi-task corpus. This evaluation focused on the base C2S-Scale checkpoints (ranging from 410 million to 27 billion parameters) that resulted from the main training phase, to measure how biological reasoning capabilities scale with model capacity prior to downstream fine-tuning.

We constructed a representative evaluation set covering the 10 pretraining tasks listed in Table 1. From the full pretraining corpus of 825 datasets, we selected 10 scRNA-seq datasets spanning a diverse range of tissues. For each of these 10 datasets, we randomly selected 5 held-out test samples for each of the 10 tasks, yielding 50 samples per dataset and a total of 500 evaluation samples. None of the cells in these test samples were seen by the model during training, ensuring that the evaluation set measures generalization to unseen cells within the training distribution.

For each C2S-Scale base checkpoint (410M, 1B, 2B, and 27B), we generated responses for all 500 evaluation samples using the latest checkpoint at each model size. We used deterministic generation with a temperature setting of 0 to remove variability in model responses. Performance metrics were calculated for each task as detailed below.

#### 4.5.1 Evaluation Metrics for Scaling Analysis

For predictive tasks such as cell type annotation and dataset interpretation, we used BERTScore [27] to measure semantic similarity between generated responses and ground truth labels. Unlike downstream fine-tuning, where models can adapt to a specific dataset’s labels and can be evaluated via exact accuracy, pretraining tasks involve diverse single-cell datasets with heterogeneous annotation schemes (e.g., varying granularity and wording of cell type labels). We therefore use BERTScore [27] as a semantic measure of cell annotation performance when evaluating base C2S-Scale checkpoints that have not received fine-tuning on any downstream datasets.

Let the reference output be *x* = ⟨*x*_1_*, …, x_k_*⟩ and the generated output be *x*^ = ⟨*x*^_1_*, …, x*^*_l_*⟩, where tokens are represented by contextual embeddings. Pairwise similarity is given by the cosine similarity 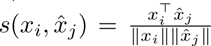 .

BERTScore recall, precision, and F1 are defined as:

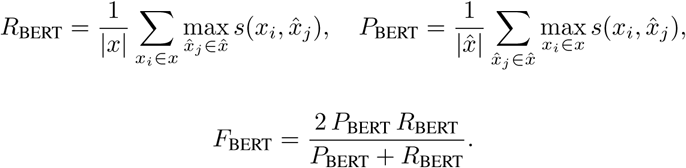

Unless otherwise noted, reported BERTScore values correspond to the F1 variant.

For generative tasks such as conditional cell generation, we evaluated outputs by measuring gene overlap between generated and target cell sentences. This metric captures the proportion of ground truth genes recovered in the generated output, defined as:

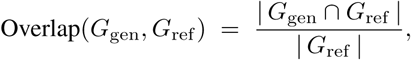

where *G*_ref_ and *G*_gen_ denote the sets of genes in the reference and generated cell sentences, respectively.

#### 4.5.2 Parameter-Efficient Scaling Analysis

To further assess scaling laws under constrained computational budgets, we conducted parameter-efficient fine-tuning experiments using Low-Rank Adaptation (LoRA) [25]. We utilized the Gemma-2 2B and 27B models as base architectures, with each training run conducted on a single NVIDIA H100 GPU. To isolate the impact of trainable parameter count, we varied the LoRA rank parameter (*r*) from 4 to 128 while holding other hyperparameters (learning rate, batch size, training data) constant. For evaluation, trained LoRA adapters were merged with their corresponding frozen base models, and performance was assessed on the same held-out multi-task test set described above.

### 4.6 Post-training methods

#### 4.6.1 Supervised fine-tuning

Following the primary training phase on the multi-task C2S-Scale corpus, base C2S checkpoints were adapted for specific downstream applications through supervised instruction fine-tuning (SFT), represented in Figure 1D (Stage 3). We used the same autoregressive next-token prediction objective employed during the main training stages (stage 1 and 2), with input prompts formatted to align with specific tasks. For example, a cell type annotation prompt appends a query instruction to an input cell sentence, triggering the generation of a metadata label for that cell.

Unless otherwise specified, we computed the loss exclusively on the response tokens during the fine-tuning phase, masking the gradients for the input cell sentences and instructions. This was done because base C2S-Scale checkpoints have already learned the transcriptomic distribution of cell sentences and the syntax of input prompts involving cell sentences during the main training phase. For fine-tuning, the optimization focus shifts entirely to refining output generation. This is particularly important for predictive tasks with short target sequences, such as cell type annotation, where a short label consisting of a few tokens contributes little to overall loss over a lengthy input context consisting of thousands of gene tokens.

For all downstream applications, we perform full finetuning to adapt all parameters for the specific task. Detailed setup and hyperparameters are provided for each downstream task in Section 4.7 below, and examples of prompt templates for each task are available in Supplementary Figures 11–16.

We note that full finetuning of base C2S-Scale models is not necessary in all cases. Parameter-efficient training strategies such as Low-Rank Adaptation (LoRA) [25] can be considered as viable alternatives for full finetuning on specific datasets of interest, providing most of the performance gain with much less computational requirements. These parameter-efficient methods update a small subset of model parameters while keeping the majority of pretrained weights frozen, allowing for rapid task-specific adaptation while requiring less computational resources.

#### 4.6.2 Reinforcement learning alignment

Reinforcement learning (RL) was used to further align model outputs with biological accuracy and interpretability. We employed Group Relative Policy Optimization (GRPO), a policy-gradient method that incorporates task-specific reward signals directly into parameter updates [50, 51].

The supervised fine-tuned model (policy *π_θ_*) generated multiple candidate outputs *o* = (*o*_1_*, …, o*_|_*_o_*_|_) for each input prompt *q*. Each token *o_t_* was assigned probability *π_θ_*(*o_t_* | *q, o_<t_*), where *o_<t_* denotes the prefix. Rewards *r_i_* were assigned to each candidate sequence *o_i_* using automated evaluation metrics such as BERTScore [27] and domain-specific scores for tasks like perturbation response prediction.

Proximal Policy Optimization (PPO) maximizes a clipped surrogate objective, which requires estimating per-token advantages *A_t_* using a value function:

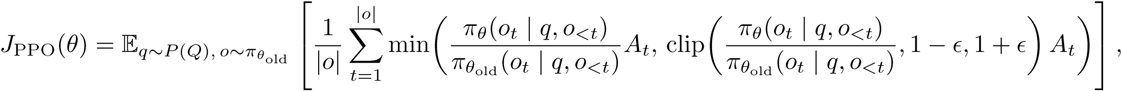

where 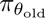 is the policy from the previous iteration, *A_t_*is the advantage at step *t*, and *ϵ* is the clipping threshold. Maintaining a critic to estimate *A_t_* increases computational cost and can destabilize training.

GRPO replaces the value function with a group-relative baseline. For each prompt *q*, the model samples *G* candidate outputs {*o*_1_*, …, o_G_*} with associated rewards {*r*_1_*, …, r_G_*}. Relative advantages are defined by normalizing rewards across the group:

The GRPO objective is

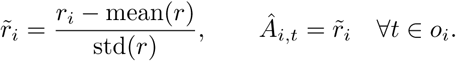

where *π*_ref_ is the frozen SFT model and *β* controls the KL regularization strength.

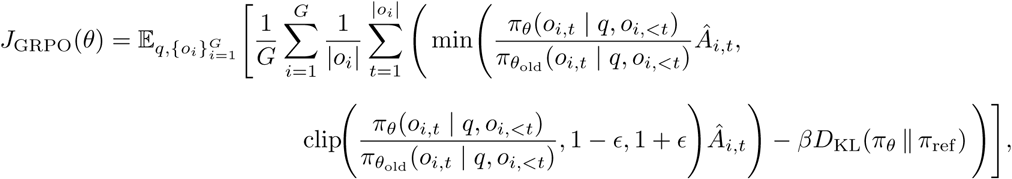

GRPO eliminates the critic network, reduces memory requirements, and yields stable optimization at scale. When trained with biologically relevant reward functions, C2S-Scale refined its predictions and aligned generative behavior with biological ground truth.

### 4.7 Downstream Tasks

#### 4.7.1 Cell type annotation

##### Datasets and Preprocessing

We benchmarked cell type annotation performance across five datasets comprising of immune, pancreas, and lung tissues. For all datasets, we performed common preprocessing steps including filtering genes expressed in fewer than 3 cells, normalizing library size to 10, 000 counts per cell, and applying a log1p transformation. The Immune 1 dataset [20] served as an unseen-cell benchmark within a scRNA-seq study that was seen during the main C2S-Scale training phase; we selected 10,000 training and 1,000 testing cells strictly from the 10% holdout cells set that was completely unseen during the main pretraining phase. All remaining datasets represented studies that were not a part of the C2S-Scale pretraining corpus, and were train/test split by patient and sample. Immune 2 [22] provided 8,000 training cells from 8 donors and 2,000 test cells from 2 held-out donors. Similarly, Immune 3 [21] also contained 8,000 training cells selected randomly from 8 donors and 2,000 test cells selected randomly from 2 held-out donors; to enable out-of-distribution (OOD) evaluation, both the Immune 1 and Immune 2 datasets were harmonized to a common set of 15 immune cell types and intersected to a shared feature space of 17,011 genes. The Pancreas dataset [23] contained 8,500 training cells from 2 donors and 3,200 test cells from 2 held-out donors. Finally, the Lung dataset [24] contained 8,892 cells from one patient for training and 6,914 cells from a held-out patient for testing.

##### Model Implementations

We compared C2S-Scale against expression-only foundation models, general-purpose LLMs, and standard single-cell classifiers. For C2S-Scale, we fine-tuned the base 1B and 2B checkpoints using the autoregressive next-token prediction objective with a learning rate of 5 × 10^−6^, weight decay of 0.0, and a batch size of 64. Input prompts combined the cell sentence with natural language instructions (Supplementary Figure 11), and predictions were evaluated via exact string matching against ground truth labels. For expression-only foundation models, we utilized official pretrained checkpoints including scGPT (human-whole-33M) [4], Geneformer (V1, 37M) [5], and scFoundation (100M) [6]. We fine-tuned the pretrained encoders with added linear classification heads using the specific training splits, utilizing official codebases for implementation and model-specific preprocessing (e.g., binning for scGPT, rank-value encoding for Geneformer). General-purpose LLMs (Gemini 1.5 Pro, GPT-4o, Llama-3.1 8B, and BioMistral 7B) were evaluated in a zero-shot setting, utilizing prompts with a task description followed by the standard C2S instruction and cell sentence. Finally, standard baselines including scVI [19], CellTypist [20], logistic regression, and PCA with a linear classifier were trained from scratch using their respective Python implementations. For linear models, input expression data was min-max scaled using the 99.9th percentile value to normalize inputs to the [0, 1] range.

##### Evaluation Protocol

All models were evaluated on the held-out test split for each dataset using classification accuracy. For the Out-of-Distribution (OOD) Generalization task, we trained all models on the training split of the Wells et al. dataset (Immune 3) and evaluated them directly on the test split of the Gong et al. dataset (Immune 2) without further adaptation. This setting tests the model’s ability to generalize annotations across different experimental batches and dataset contexts while maintaining consistent cell type semantics.

#### 4.7.2 Cell generation

##### Task Setup and Evaluation

Unlike downstream tasks such as perturbation prediction (detailed in Section 4.7.9), the cell generation task serves primarily as an evaluation of one the model’s fundamental pretraining tasks: the ability to synthesize biologically coherent gene expression profiles from natural language instructions. We evaluated base C2S-Scale checkpoints immediately following the main training phase on the held-out evaluation set described in Section 4.5.

For each test sample, the model was provided with a natural language prompt containing relevant metadata (e.g., “Generate a list of genes… representing a Homo sapiens cell of cell type B cell”). See Supplementary figures 12 for examples. The model was then tasked with generating a valid cell sentence (or multiple sentences) based on the metadata, which involves determining the expressed genes and their rank order. We utilized deterministic decoding (temperature = 0.0) for all quantitative benchmarking. We note that while higher temperatures (*T >* 0) are generally preferred for recovering the full distributional diversity of scRNA-seq data in generation tasks, deterministic generation provides more reproducibility and reduces variance in the model’s outputs.

##### Metrics

Generated cell quality was quantified using the **Expressed Gene Overlap Percentage**, defined in Section 4.5. This metric calculates the set intersection between the genes present in the model-generated cell sentence and those in the corresponding ground-truth cell sentence.

We acknowledge that this metric is conservative and may penalize the model for dataset-level shifts. Since the input prompts are often dataset-agnostic (specifying only tissue or cell type), the model may validly generate a B cell profile resembling a dataset with a different gene panel or batch effect than the specific ground-truth cell used for comparison. Despite this penalization, we found gene overlap to be a reliable proxy for the model’s generation ability when given biological metadata constraints. For more fine-grained generative evaluations accounting for distributional matching, we refer to the Perturbation Prediction methods (Section 4.7.9).

#### 4.7.3 Cell embedding

##### Task Setup and Datasets

To evaluate the quality of learned representations without task-specific fine-tuning, we benchmarked models on the **Immune 1**, **Pancreas**, and **Lung** datasets, utilizing the identical train/test splits described in Section 4.7.1.

##### Embedding Extraction

We extracted frozen latent representations from three distinct model categories. For **C2S-Scale**, we utilized the base pretrained checkpoints (e.g., C2S-Scale 1B) with input prompts formatted identically to the cell type prediction task. Instead of autoregressive generation, we performed a single forward pass and extracted the hidden states from the final transformer layer, applying average pooling across all tokens in the sequence (excluding padding) to obtain a single fixed-length vector per cell. For **expression-only foundation models**, we evaluated the same pretrained checkpoints used in the annotation task (scGPT-whole-human-33M, Geneformer V1-37M, scFoundation-100M). We used their respective official inference pipelines and preprocessing schemes to extract latent representations, applying average pooling to the final layer embeddings to ensure consistency with the C2S-Scale methodology. Finally, for **general-purpose LLMs**, which cannot process raw transcriptomic counts, we utilized the text embedding APIs provided by OpenAI (for GPT models) and Google (for Gemini models) to generate embeddings for cell sentence text representations. While these models are not explicitly trained on biological data, this approach assesses the extent to which general natural language pretraining captures semantic structure within cell sentences.

##### Evaluation Protocol

We assessed embedding quality using a standard linear probing protocol. For all embedding models (C2S-Scale, expression FMs, and general LLMs), we trained a regularized linear classifier (Logistic Regression) on the embeddings of the training set and evaluated classification accuracy on the test set embeddings. For simple baseline models (scVI, CellTypist, PCA), which do not produce generalized pretrained embeddings in the same manner, we report their supervised classification performance from the **Cell type annotation** section as a reference benchmark.

This comparison serves to highlight the trade-off between task-specific training (simple baselines) and generalized representation learning (foundation models).

#### 4.7.4 Single-cell bulk integration

Multimodal integration is essential for capturing the complexity of biological systems, as different data modalities provide complementary perspectives on cellular function. Each modality has its own strengths and limitations: some offer high resolution at the cost of sparsity, while others provide broader coverage but lack single-cell detail. Therefore, models that can integrate modalities can provide a more complete and robust understanding of cellular behavior, improving both interpretability and predictive power in biological analysis.

To assess this, we designed a simple single-cell and bulk RNA seq integration task. Using a single-cell lung tissue dataset [24], we constructed pseudo-bulk samples by aggregating over donor, cell type, and batch. For each pseudo-bulk sample, we randomly sampled ten single-cell samples from the same conditions to construct pairs. We embedded each single-cell and pseudo-bulk sample individually using each model and computed the cosine similarity between the paired single-cell and bulk samples. Following [66], we used the “fraction of samples closer than the true match” (FOSCTTM) to evaluate the performance of each model. A FOSCTTM of 0 corresponds to a perfect model (the cosine similarity of matched pairs is higher than any other pair), whereas a FOSCTTM close to 0.5 means the cosine similarity between the matched pairs is about as good as the cosine similarity between random pairs.

#### 4.7.5 Cluster captioning

##### Dataset Construction

To generate a high-quality ground truth dataset for cluster-level interpretation, we curated 30 scRNA-seq datasets and performed standard preprocessing, clustering, and differential expression analysis. We utilized GPT-4o [17] to generate descriptive captions for each cluster; to ensure biological accuracy, the generation prompt included the cell type, tissue, organism, disease state, top three differentially expressed genes, and the full text of the associated publication. This process yielded 1,723 captions spanning 345 distinct clusters. For each training sample, we randomly sampled two cells from a cluster to form a multi-cell context prompt (Supplementary Figure 14), with the natural language caption serving as the target output.

##### Task Setup and Fine-tuning

Cluster captioning serves as a specific downstream task distinct from the main multi-task training phase. We fine-tuned base C2S-Scale checkpoints using standard supervised fine-tuning (SFT) with a learning rate of 1 × 10^−5^, weight decay of 0.01, and a batch size of 64. Consistent with our general post-training protocol, training utilized the next-token prediction objective with the loss calculated exclusively on the caption tokens.

##### Baseline Models

We compared C2S-Scale against two categories of models capable of natural language generation. First, since standard expression-only foundation models (e.g., scGPT) lack text decoding capabilities, we benchmarked against General-purpose LLMs (GPT-4o and Gemini 1.5 Pro) in a zero-shot setting. These models received the identical multi-cell sentence inputs and instructions used for C2S-Scale. Second, we benchmarked against CellWhisperer [67], a specialized multimodal model designed for cluster annotation. Unlike C2S-Scale, which processes sequences of individual cells, CellWhisperer operates on aggregated (pseudo-bulk) gene expression profiles via an encoder aligned with a text decoder. We utilized the official pretrained CellWhisperer model and inference pipeline for comparison.

##### Evaluation

All models were evaluated on a held-out test set of clusters that were strictly unseen during C2S-Scale fine-tuning. Performance was quantified using BioBERTScore [27], measuring the semantic similarity between the model-generated caption and the ground-truth description derived from the study context.

#### 4.7.6 Dataset interpretation

##### Task Setup and Test Set Construction

To assess the model’s ability to produce high-level biological insights from raw transcriptomic data, we designed a dataset-level natural language interpretation task. Models were presented with a multi-cell context (5-20 cells sampled from the same tissue and donor) and prompted to generate a biological abstract summarizing the experiment. To prevent memorization of the original study abstracts, we generated 500 variations of each ground-truth abstract using GPT-3.5-Turbo, increasing diversity in language and phrasing while preserving factual content (Supplementary Figure 13). Since dataset interpretation was included as a main stage training task (Table 1), we evaluated base C2S-Scale checkpoints immediately following the main training phase without additional fine-tuning. We constructed two separate evaluation sets to measure performance. The In-Distribution (ID) Test Set comprises 3,065 samples derived from 613 scRNA-seq datasets within the C2S-Scale pretraining corpus; while these specific samples were strictly held out from training, the model had been exposed to other cells and abstract samples from these same studies. To rigorously benchmark generalization to unseen biological contexts, we constructed an Out-of-Distribution (OOD) Test Set comprising 400 samples from two studies completely unseen during pretraining: a pancreas dataset [23] and a human retina study [68].

##### Baseline Models and Evaluation

Expression-only foundation models such as scGPT, Geneformer, and scFoundation were excluded from this benchmark as they lack the capability to process natural language instructions or generate textual summaries. Instead, we benchmarked against state-of-the-art general-purpose LLMs including Llama-3, GPT-4o, and Gemini 1.5 Pro. These baselines received identical inputs to C2S-Scale: a system prompt defining the task, followed by the raw cell sentence representations of the multi-cell context. This comparison isolates the value of domain-specific pretraining on transcriptomic sequences against the general reasoning capabilities of frontier LLMs. Given the open-ended nature of abstract generation, exact string matching is insufficient for evaluation. We therefore quantified performance using BERTScore (F1) [27], measuring the semantic similarity between the model-generated summary and the ground-truth abstract variations. This metric was selected for its superior correlation with human judgment compared to n-gram based metrics like ROUGE or BLEU.

#### 4.7.7 Spatial niche prediction

For the spatial niche prediction task, we used the CosMx Spatial Molecular Imager Human Liver dataset [39], which provides annotated spatially-resolved single-cell data from both normal and hepatocellular carcinoma liver tissues from two different donors. This dataset encompasses over 800,000 single cells across a total of approximately 180 mm^2^ of liver tissue, with expression measured on a set of 1,000 curated genes. The dataset was processed to filter out genes expressed in fewer than three cells and cells expressing fewer than 50 genes. It was then normalized to a total count of 1 × 10^4^ and the base 10 logarithm was applied. Spatial coordinates were saved to define neighborhoods and facilitate spatial analyses. We define a neighborhood to be a radius of 0.02 pixels (approximately 20 *µ*m), chosen to maximize the number of cells we can fit into the model’s context. The dataset was split into training and test sets based on spatial coordinates to prevent spatial leakage between sets.

To train C2S-Scale on spatial and multi-cellular relationships, we designed the following tasks. **1. Spatial neighborhood prediction:** Given multiple cell sentences, predict whether these cells come from the same neighborhood. **2. Conditional neighbor generation:** Given multiple cell sentences from a neighborhood, generate a novel cell sentence that would belong to the same neighborhood. **3. Niche label prediction:** Given a cell sentence for a single cell, predict the niche label annotation for that cell. **4. Same niche prediction:** Given multiple cell sentences, predict whether all of these cells have the same niche label or different niches.

To construct prompts, cell sentences were randomly sampled from the appropriate data split. Multi-cell contexts were created by taking all cells in the sampled cell’s neighborhood for positive samples, or an equivalent number of randomly sampled cells outside the neighborhood as negative samples. The data contained 19,754 training samples and 3,968 test samples. Example formatted prompts for spatial tasks can be found in Supplementary Figure 16.

Additionally, to enhance the model’s understanding of cell communication, we included gene interaction metadata from CellPhoneDB [41] and BioGRID [42] in the training data mixture. We restricted the data to only retain interactions involving the 1,000 genes in the CosMx data, and also only to genes coding for extracellular proteins (determined using MatrixDB [69]). We included 5,822 interaction samples from CPDB and 2,334 from BioGRID.

Models were evaluated on a held-out test set comprising 3,968 samples. Performance was quantified using binary classification accuracy for the predictive tasks, and the gene percent overlap (Jaccard index) for the neighbor generation task. C2S models and GPT-4o were given the natural language prompts consisting of cell sentences and were asked to predict the output in natural language. As scGPT and Nicheformer cannot natively handle multicellular inputs, we averaged the embeddings for each cell in the input and fit a linear classifier on the averaged embeddings. The “Linear Model” baseline was fit on the average expression vector.

To compare models for the predictive tasks, paired differences in prediction outcomes were assessed using McNemar’s test with continuity correction, which evaluates whether two classifiers differ significantly in their error distributions when applied to the same test set. For the generative task, we generate 100 replicates for each test, and compute significance using the two-tailed t-test. Significance was reported as p-values from the corresponding test, with values below 0.05 considered statistically significant.

#### 4.7.8 Question answering

##### Dataset Construction

To assess the model’s ability to reason jointly over transcriptomic data and natural language, we constructed a multimodal QA dataset. We utilized GPT-4.5 to generate high-quality question-answer pairs combining cell sentences and specific biological questions about multi-cell samples from scRNA-seq datasets. To create the QA pairs, GPT-4.5 was provided with a hybrid context consisting of: (1) key text sections (Abstract, Results, Discussion) from a published scRNA-seq study, and (2) raw cell sentences sampled from the corresponding dataset. GPT-4.5 was tasked with formulating biologically meaningful questions that required synthesizing information from both the gene expression profiles and the textual findings (e.g., relating specific marker gene expression in the cell sentence to cell type definitions in the text). This process yielded approximately 1,600 QA pairs across 80 diverse studies (20 QA pairs per study).

##### Two-Stage Training Pipeline

We fine-tuned C2S-Scale for question answering using a two-stage approach:

1. **Supervised Fine-Tuning (SFT):** We first fine-tuned base C2S-Scale checkpoints using standard SFT. The model was trained to generate the target answer given the question and cell sentence context. We used a learning rate of 1 × 10^−5^, a global batch size of 64, and 100 linear warmup steps for fine-tuning.
2. **Group Relative Policy Optimization (GRPO):** To further align model outputs with ground truth answers, we applied GRPO following SFT. We constructed a separate RL training set of 600 samples from unseen studies. During GRPO training, the SFT model generated 32 candidate answers for each prompt. We calculated a reward signal for each candidate using **BioBERTScore**, which measures the semantic alignment between the candidate and the ground truth answer. This reward guided the policy update to favor answers that were factually accurate and biologically context-aware. For GRPO, we set the KL-divergence coefficient *β* = 0.03 and used a learning rate of 5 × 10^−7^.

##### Baselines and Evaluation

We benchmarked C2S-Scale against state-of-the-art general-purpose LLMs, including GPT-4o and Gemini 1.5 Pro. Standard single-cell foundation models (e.g., scGPT, Geneformer) were excluded from this comparison as they lack the natural language decoding capabilities required for open-ended question answering. All models were evaluated on a held-out test set derived from studies completely unseen during both the SFT and GRPO phases. Performance was quantified using BERTScore to assess the semantic quality of the generated answers.

#### 4.7.9 Perturbation prediction

##### Cytokine Stimulation

The Dong et al. dataset [53] dataset includes immune cells exposed to individual and combinatorial cytokines, with each cell annotated by type, stimulation, and exposure length – yielding 133 conditions. We retained the 5000 most highly variable genes to prioritize high-signal features and align with established benchmarks in perturbation modeling, which typically utilize between 1000 and 7000 genes [4, 70, 71, 72, 73], and evaluated models in the scGPT embedding space [4] using maximum mean discrepancy (MMD), Wasserstein distance, and scFID (Section 4.8). This embedding-based evaluation provides more meaningful comparisons than expression-level metrics, which can be skewed by a small number of genes with extreme values.

The training of C2S models for the Dong et al. dataset followed a structured two-stage process to effectively predict responses to unseen cytokine stimulations. The test dataset featured three tiers of held-out perturbations with increasing difficulty: (1) a completely excluded combinatorial perturbation (interferon-*β* + IL-6), (2) one perturbation entirely held out for each cell type across both chronic and acute conditions (B: interferon-III, CD4 T: interferon-*γ*, CD8 T: interferon-*α*2, Dendritic: interferon-*β* (no chronic cells), NK: IL-6), and (3) one perturbation excluded in either chronic or acute conditions for each cell type while the other condition remained in training (B: acute interferon-*β*, CD4 T: acute interferon-*β* + interferon-*γ*, CD8 T: chronic TNF-*α*, NK: chronic interferon-III). In the first stage, the model was fine-tuned using supervised learning on both cell sentence generation and natural language label prediction, where it simultaneously predicted all three labels—cell type, perturbation, and exposure—ensuring it learned bidirectional relationships between conditions and gene expression. This fine-tuning stage was conducted for 3–4 epochs using the Hugging Face Trainer on a single H100 GPU. Example prompts for training can be found in Supplementary Figure 15, PBMC.

The second stage employed GRPO to refine perturbation response generation. For the Dong et al. dataset, the reward was computed as the negative mean squared error between generated and ground truth cells, randomly paired under the same condition labels and embedded using scGPT. GRPO training used 32 generated responses and 32 real cells per prompt, and was conducted on 4 H100 GPUs for 3 epochs. The interferon subset used for GRPO was defined as the union of the MSigDB [74] interferon-*α* and interferon-*γ* hallmark gene sets, intersected with the highly variable genes (HVGs) from the dataset, resulting in 95 genes.

To benchmark against other perturbation response models, we included scGen, BioLord, CellOT, CondOT, scGPT, and linear baselines. For scGen [72], we used the pertpy library [75] to train and run inference. In order to generate meaningful perturbations for unseen combinatorial perturbations, we adapt the model by summing the learned latent stimulated vectors from each individual cytokine perturbation to gather the latent stimulated vector required for scGen’s predict, rather than treating combinatorial perturbations as a separate condition altogether. BioLord [71], which is built on top of scGen’s infrastructure, naturally handles unseen combinations of conditions as long as the individual conditions are seen during training in some other combination. To this end, we treated each cytokine perturbation as a combination of two attributes to allow for the possibility of two cytokine stimulations appearing in combination, which natrually fits into BioLord’s framework. For CellOT, we followed the training procedure described in [73] using a pre-trained scGen autoencoder on all training data and a feed-forward network for transport. In order to allow for generalization to unseen treatments needed for the benchmark, we train a single transport map per cell type and use learned embedding layers for one-hot representations of individual cytokine treatments and exposure values. For combinatorial perturbations, the embeddings of each cytokine are summed together. Then, the embedded cytokine and exposure vectors are summed with the latent encoding of the cell prior to transport. In contrast, we do not make any modifications to the architecture of CondOT [76], which is naturally suited for unseen conditions and trains a unified model for all condition types. For scGPT [4], we added linear encoders for cell type, perturbation, and exposure, projecting binary vectors into dense vectors, and then added these embeddings to each gene token embedding before forwarding them through the model. This is a minimal modification compared to the original genetic perturbation scGPT model, which instead provides a binary signal to each gene indicating whether it is affected. Out of the 5000 selected highly variable genes, 3801 were in scGPT’s pre-trained gene vocabulary. Since adding new tokens can significantly reduce a pre-trained transformer model’s performance, we only trained and evaluated scGPT on this subset of genes. Finally, our linear baseline models are ridge regression models with binary encodings for conditions. The PCA variant predicts the gene expression values directly from the 100 principal components plus the same binary encodings for conditions. We found this model performed better than making predictions in PCA space and applying the inverse transform.

##### Small Molecule Perturbations in Bulk RNAseq

For the L1000 dataset [52], we trained on the 978 genes measured in the original study, referred to as *landmark genes*, using the level 3a data, which normalizes the measured genes with fitted invariant gene set curves and quantile normalization (see [52] for further details). We paired untreated and treated samples by matching the cell line name. To evaluate generalization, we selected 50 perturbations with fewer than 1,000 total samples and held out half the cell lines in each perturbation as test data. We used Kendall’s *τ* as the reward function during reinforcement learning, as it properly accounts for tied ranks. This is especially important for L1000 where non-expressed genes share the same lowest rank. SFT was conducted using a batch size of 2 and gradient accumulation of 32, with a learning rate of 1e-4. Training ran on a single H100 GPU for 3,500 steps (approximately one epoch, though not all data is seen due to dataset size). For GRPO, the model was trained with a batch size of 8 and gradient accumulation of 4. We generated 24 responses per prompt. The learning rate was set to 1e-6 with a beta value of 5e-3. Training was distributed across 4 H100 GPUs—three for model training and one for vLLM-based response generation. GRPO ran for approximately 3,000 steps over 3 epochs, although as with SFT, the model likely saw less than a full epoch due to data scale. Example prompts for training can be found in Supplementary Figure 15, L1000.

For evaluation, we computed metrics differently across datasets. For the Dong et al. [53] dataset, we computed maximum mean discrepancy (MMD), Wasserstein distance, and scFID for each unique combination of condition labels (cell type, cytokine, and exposure duration), and averaged these values across all combinations to obtain the final metric. For the L1000 dataset [52], we computed Pearson’s *r* against the Level 3 gene expression values and Kendall’s *τ* on the ranks of the gene expression values for each test sample individually and then reported the average across all samples.

Kendall’s *τ* measures rank correlation between two ordered lists. Given *n* genes, we consider all ^1^ *n*(*n* − 1) possible gene pairs. For any pair of genes (*i, j*), if their relative order (which gene is ranked higher) is the same in both the generated output and the ground-truth ranking, the pair is *concordant*; if their relative order is reversed, the pair is *discordant*. Tied pairs (where the genes share the same rank in either list) are handled by assigning them the same value. Kendall’s *τ* is then defined as

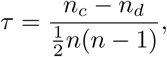

where *n_c_* and *n_d_*denote the number of concordant and discordant pairs, respectively. In our application, the ranks of the 978 L1000 landmark genes are derived from the generated output of the model, where the cell sentence places genes in descending expression order (e.g., GeneT GeneA GeneS GeneW …). Genes not present in the model’s output are assumed to share the lowest possible rank (e.g., if 950 genes are generated, the remaining 28 share rank 951). The same ranking convention is applied to the L1000 ground-truth sample, where unexpressed genes also share the last rank. Kendall’s *τ* is then computed between these two ranked lists, yielding a rank-based correlation that is robust to tied ranks and sparse expression. Only the apoptosis genes from the MSigDB hallmark set that were present in the L1000 landmark gene list were used during GRPO, totaling 40 genes.

##### Small Molecule Perturbations in single-cell RNAseq

We utilized the SciPlex3 dataset [55], which consists of 187 small molecule perturbations applied across three cell lines (A549’, MCF7’, ‘K562’). Our training and evaluation followed the out-of-distribution setup described in ChemCPA [56], where the test set consists of 9 drugs completely unseen during training: Dacinostat, Givinostat, Belinostat, Hesperadin, Quisinostat, Alvespimycin, Tanespimycin, TAK-901, and Flavopiridol. This split results in a training set of 501,004 cells (excluding validation cells) and a test set of 22,777 cells. We employed the 2,000-gene feature set established in ChemCPA, comprising the 978 landmark genes plus the 1,022 highest variable genes not present in the landmark set.

We trained each C2S model on 4 H100 GPUs with an effective batch size of 128 and a learning rate of 1e-4. The 410M parameter model was trained for approximately 12 epochs, and the 1B parameter model for approximately 8 epochs, at which point the validation loss converged. To limit token usage and sequence length, C2S models utilized pathway information contained in the SciPlex data rather than RDKit embeddings. Example prompts for training can be found in Supplementary Figure 15, SciPlex.

We benchmarked against ChemCPA, scGPT, and CondOT. We used the ChemCPA model with the autoencoder architecture pretrained on L1000 data, as this configuration yielded the best performance in the original study. To allow for the generation of unseen perturbations and ensure fairness regarding molecule data access, scGPT was adapted to use RDKit [77] embeddings; similar to the cytokine setup, we added cell type and dose embeddings along with a linear projection of the RDKit embeddings. CondOT was similarly adapted to use RDKit [77] embeddings for the same reasons. For evaluation, we generated the same number of cells for each unique combination of cell line, dose, and drug as present in the test set. We computed the same metrics as in the cytokine stimulation task (MMD, Wasserstein distance, and scFID) for each combination and reported the average across all combinations.

### 4.8 Single-Cell Fréchet Inception Distance

The scFID is an adaptation of the FID [54] tailored for evaluating generative models in single-cell transcriptomics. While the traditional FID employs the Inception v3 model [78] to extract features from images, scFID utilizes scGPT [4] as its foundation model to embed single-cell gene expression profiles. Notably, scFID is flexible and can incorporate any suitable foundation model for embedding. The scFID quantifies the similarity between the distributions of real and generated single-cell embeddings by assuming that these distributions are multivariate normal (Gaussian). Under this assumption, the scFID computes the Wasserstein distance between the two Gaussian distributions, providing a measure of how closely the generated data resembles the real data in the embedding space.

Mathematically, given two sets of single-cell embeddings—one from real cells and one from generated cells—scFID is defined as:

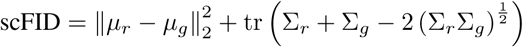

where:

- *µ_r_* and *µ_g_* are the mean vectors of the real and generated cell embeddings, respectively,
- Σ*_r_* and Σ*_g_* are the covariance matrices of the real and generated cell embeddings, respectively,
- tr denotes the trace of a matrix.

To evaluate generative model performance across various conditions, we computed the scFID for each unique combination of test labels—such as specific cell types, perturbations, and exposure durations—and then averaged these individual scFID values.

To validate that scFID accurately reflects biological similarity within the foundation model’s embedding space, we performed a control analysis using the Dominguez immune dataset [20]. We hypothesized that the distributional distance between subsets of cells sharing the same biological state should be minimal compared to distances between distinct states. As shown in Supplementary Figure 19, scFID values computed within identical cell types were significantly lower than those computed between different cell types (two-sample t-test, *P* = 2.23 × 10^−4^). This distinction was even more pronounced when analyzing unsupervised Leiden clusters (two-sample t-test, *P* = 2.60 × 10^−9^). These results indicate that the metric effectively discriminates between biological distributions in the latent space, supporting its utility as a measure of generative quality.

### 4.9 Virtual Screen Setup

#### Perturbation Prediction Model and Delta Sentences

For high-throughput virtual screening, we utilized a specialized C2S-Scale perturbation model initialized from a Gemma-3 4B checkpoint [79]. This model was trained on a perturbation corpus combining the LINCS L1000 bulk RNA-seq dataset [52] and the Tahoe-100M single-cell dataset [80], spanning thousands of drug perturbations across hundreds of cell lines. To maximize computational efficiency for large-scale screening, we employed a *delta sentence* prediction objective rather than generating the entire cell sentence of the target perturbed profile. A delta sentence captures the vector difference between a perturbed and control state (Δ*E* = *E*_stimulated_ − *E*_control_). The model was trained to output the top 200 upregulated and top 200 downregulated differentially expressed genes (DEGs) formatted as two newline-separated, rank-ordered lists. During inference, these predicted delta sentences are mapped back to quantitative log-fold changes using pre-fitted log-linear models and added to the input control profile to yield the final predicted transcriptome.

#### Datasets

We analyzed drug responses in both primary tumor samples and an immortalized cell line in order to capture effects across distinct immune environments. The immune-context-positive data comprised flow sorted bulk RNA-seq from a pan-cancer atlas [81], which includes 364 tumor specimens spanning 12 cancer types. We restricted our analysis to cells labeled as “tumor cells” from this dataset, yielding 162 bulk samples. As an immune context-neutral system, we used data from GEO for the WAGA (Merkel cell; GSE130346), MCC2 (Merkel cell; GSE223275 [82]), and UTSCC45 (squamous cell; GSE122512 [83]) cell lines, as they are not part of the training data for the model. For the single-cell data, standard preprocessing was applied, including removal of genes expressed in fewer than three cells, removal of cells with fewer than 50 counts, normalization to a total count of 10^4^ per cell, and log1p transformation.

To quantify type I interferon activity across contexts, we computed the gene set score for the “Hallmark Interferon Alpha Response” gene set (genes listed in Supplementary Fig 20) using Scanpy’s score_genes function (version 1.9.8).

#### Compound library

The screening library was derived from the L1000 resource, which catalogs over 30,000 small molecules. We first manually filtered down the full library of drugs to approximately 4,000 drugs that had common names. We then searched through the remaining drugs which were represented with BRD-XXXXX identifiers specific to the LINCS library, to see if any were common drugs that had vendors. Using the Repurposing Data Portal from the Broad Institute as well as PubChem vendor matching, 242 of the BRD drugs were found to have common names or vendor information, and were added to our screening library for experiments. In a few cases, o3 deep research was used to double-check that vendors were available for named drugs. This produced a final working library of 4,266 drugs, which was used for virtual screening experiments.

#### Perturbation inference

Drug perturbations were simulated for all drugs (N=4,266) using our C2S-Scale perturbation response prediction model. Each bulk tumor sample was perturbed *in silico* with every drug in the library three times, for a total of *N* = 486 samples per drug. For the WAGA cell line, 20 representative cells were each perturbed 20 times with every drug for a total of *N* = 400 samples per drug. Replicates corresponded to independent forward passes through the model, with stochastic sampling at a temperature of 0.3 to introduce variability across predictions.

#### Scoring of antigen-presentation programs

Antigen-presentation activity was quantified by calculating enrichment scores for each perturbed profile. We applied single-sample gene set enrichment analysis (ssGSEA) with the “Class I MHC mediated antigen processing and presentation” gene set from MSigDB [74], using the Python package gseapy (v1.1.8) with parameters sample_norm_method=‘rank’ and weight=0. The genes contained in this gene set are listed in Supplementary Fig. 21. Scores were aggregated across replicates for each drug and z-scored.

Top-ranked drugs (score *>* 0.75, *P <* 0.05) were examined for prior evidence of involvement in antigen-presentation pathways using Google Gemini, and reviewed by two oncologists (Supplementary Table 3). Manual inspection was used to flag positive controls and compounds not previously reported in the literature, and these were prioritized for further analysis.

### 4.10 Experimental Validation in Tumor Fragment and Cell Line Models

#### 4.10.1 Ethical approval

The study was approved by the Institutional Review Boards at Yale University following Yale Spore in Skin Cancer (IRB protocol #0609001869). Patients with a clinical indication for surgical resection consented to the donation of surgically resected tissue for research use.

#### 4.10.2 Primary tumor modeling

Patient-derived melanoma organoids (PDMOs) were established from excess surgical specimens obtained with informed consent. Fresh tumor tissue was minced on ice and enzymatically digested (collagenase/dispase plus DNase I) to generate a single-cell/small-cluster suspension, passed through a 100-*µ*m strainer, and washed in ice-cold basal medium. Cell pellets were resuspended in growth factor–reduced Matrigel and plated into pre-warmed 24-well plates, then overlaid with melanoma organoid medium (Advanced DMEM/F12 supplemented with B27, N2, EGF, FGF, Noggin, R-spondin, and ROCK inhibitor for the first 48 h). Recombinant IL-2 was added to support the survival of resident immune cells and maintain autologous tumor–immune interactions during early organoid formation. Organoids were maintained at 37^◦^C and 5% CO_2_ for 48 h up to one week, with medium changes every 2–3 days, during which time they adopted characteristic organoid morphology and melanoma organoids developed visible pigment under phase-contrast microscopy. For drug validation, mature organoids were gently dissociated into small fragments (≈50–100 *µ*m), counted, and seeded into 96-well plates in organoid medium. After overnight recovery, organoids were treated with targeted agents (crizotinib, afatinib, GSK-126, silmitasertib, trametinib at 10 nM (all SelleckChem), or 10 IU/ml IFN-*γ* (Peptrotech) in serial dilutions for 48 h, with DMSO as vehicle control. For selected conditions, parallel wells were harvested, dissociated to single cells, and stained for surface HLA-A/B/C and lineage markers followed by flow cytometry to correlate drug response with changes in antigen-presentation phenotype.

#### 4.10.3 Cell line modeling

To validate the interferon-conditional effects predicted in silico, we performed experiments in two tumor-derived cell lines: A375 (BRAF-mutant melanoma), MDK-wildtype WAGA (Merkel cell carcinoma), and MDK-knockout WAGA. Cells (600,000–2,500,000 cells/ml) were treated with either trametinib (A375) or silmitasertib (MDK-WT, MDK-KO WAGA) at the indicated concentrations for 24 hours, followed by stimulation with 2 U/ml human IFN*β* (PBL Assay Science, cat. #11415) or 2 U/ml human IFN*γ* (PBL Assay Science, cat. #11500) for an additional 24 hours. In parallel, dose–response assays were performed by titrating IFN*β* across a range of 0.5–200 U/ml to characterize sensitivity to interferon signaling.

After treatment, cells were harvested and stained for surface expression of major histocompatibility complex class I molecules HLA-A,B,C (clone W6/32, BioLegend). Live tumor cells were gated using Zombie Aqua fixable viability dye (BioLegend) to exclude dead cells prior to analysis by flow cytometry using the CytoFLEX S running CytExpert 2.4 (all Beckman Coulter). All assays were performed in three independent biological replicates. For statistical comparisons, a two-way Student’s t-test was applied, followed by the Benjamini-Hochberg correction for multiple testing.

### 4.11 Data Availability

A list of HCA and CELLxGENE datasets used for pretraining is provided in Supplementary Table 2. Spatial transcriptomic data for the niche prediction task was obtained from CosMx [39]. Publicly available interaction databases were acquired from [41, 42, 69]. For the perturbation prediction task we used transcriptomic data from L1000 [52] and from [53]. For the virtual screen we used primary tumor data from [81] and cell line data from [84]. Model weights are available on Hugging Face.

### 4.12 Code Availability

Code for model training is publicly available at: https://github.com/vandijklab/cell2sentence

## 5 Acknowledgements

The authors thank collaborators and contributors from across institutions for their invaluable support and insights throughout this project. This work was supported in part by the National Institutes of Health (NIH) grant R35GM143072–01 and the Yale Colton Center Award, both awarded to Dr. David van Dijk. BioRender (https://BioRender.com) was used for Figures 4, 5, and 6.

## 6 Author Contributions

Conceptualization, overall vision, project leadership, and supervision were done by D.v.D. Project lead and coordination were done by S.A. Rizvi. Model training was done by D.L., A.P., S.Z., E.W., and B.P. Scaling evaluations were done by S.A.R. Benchmarking across predictive and generative single-cell tasks was done by S.A.R. and A.P. Perturbation prediction evaluations and scFID implementation were done by D.L. Question answering evaluations were done by S.Z. Virtual screening was done by A.P, S.A.R, and E.W. Wet-lab experiment conceptualization was done by J.I. Wet-lab validation experiments were done by C.J.P., N.M.C., and Z.F. Data curation was done by S.H., D.Z., Z.L., C.L., E.S., D.J., and L.Z. Writing of the original draft was done by S.A.R., D.L., A.P., S.Z., and D.v.D. Writing, review, and editing were done by all authors.

## 7 Supplementary

### 7.1 FLOPs Usage and Computational Comparison

#### 7.1.1 Computational Footprint and Scaling Analysis

To ensure transparency and facilitate future research, we detail the computational resources required for training the C2S-Scale model family. We have calculated the total Floating Point Operations (FLOPs) for the primary pretraining phase across all C2S-Scale models and report these, alongside the FLOPs of comparator models, in Supplementary Table 1.

As C2S-Scale is a multimodal foundation model designed to process and reason over both large-scale transcriptomic data and rich textual corpora, its pretraining utilizes a greater number of FLOPs compared to specialized, expression-only foundation models. For instance, the C2S-Scale-Gemma-2 2B model uses a higher overall FLOP count than scGPT. This increased compute is fundamental to C2S-Scale’s ability to span an unprecedented range of single-cell tasks—including perturbation prediction, cell-type annotation, and complex natural language interpretation—all within a single, unified architecture. The resulting performance gains, which scale consistently with increasing FLOPs and model size (Figure 2), validate this strategic investment in a comprehensive, multimodal foundation model. We show that the superior performance of C2S-Scale across the entire spectrum of prediction, generation, and reasoning tasks is a direct consequence of scaling both the model size and the complexity of the integrated (transcriptomic and textual) training data, positioning it as a robust next-generation platform for biological discovery.

### 7.2 Limitations

#### 7.2.1 Addressing Limitations of Causal Attention in Gene Expression Modeling

While our approach demonstrates strong empirical performance in modeling single-cell gene expression using autoregressive language models, we acknowledge that causal attention’s inherent unidirectionality—favoring high-to-low gene expression dependencies—could theoretically limit the modeling of true causal biological interactions that flow from low- to high-expression genes. However, we contend that this constraint does not significantly impede our objectives and can be mitigated through several complementary strategies. First, our approach aligns with successful paradigms from vision-language models, where arbitrary tokenization orders paired with causal attention still achieve state-of-the-art performance [85]. Similar to hybrid vision architectures that combine causal and non-causal attention layers, our framework could incorporate indirect bidirectional context through auxiliary reasoning tokens or non-causal gene interactions.

##### Multi-cell context and reasoning as a corrective mechanism

The model’s reasoning capabilities provide additional corrective potential. Emerging evidence from language modeling demonstrates that explicit reasoning steps can compensate for causal attention limitations [86, 87, 88]. In our context, intermediate tokens representing biological pathways or gene interactions enable iterative prediction refinement, effectively circumventing strict unidirectionality. Furthermore, our multi-cell training framework enables implicit bidirectionality—low-expression genes in one cell can influence high-expression genes in the following cell, approximating bidirectional attention across a multi-cell context.

##### Correlation, not causation

It is important to emphasize that our model is designed to capture predictive correlations over inferring causal gene relationships. This mirrors natural language processing, where autoregressive models successfully capture statistical correlations despite occasional misalignment between word order and true causal relationships (e.g. passive constructions) [89, 90]. Our results confirm that expression correlations provide sufficient predictive power for key biological analysis tasks.

##### Architectural enhancements

Looking forward, we propose three architectural enhancements to further mitigate this limitation: (1) bidirectional attention by partitioning gene sequences, (2) variable gene ordering during training to induce order invariance, and (3) hybrid attention architectures blending causal and non-causal attention layers. While our current approach already demonstrates that sequential modeling of gene expression—despite lacking natural ordering—leverages pretrained LLMs without requiring custom architectures, these enhancements aim to further improve biological fidelity and predictive power.

In summary, while causal attention restricts bidirectionality within individual cells, its ability to capture correlations aligns with our predictive objectives. The combined effects of multi-cell context, reasoning mechanisms, and prospective architectural improvements position this approach as a robust foundation for single-cell analysis, with multiple pathways available for extending its biological fidelity.

#### 7.2.2 Hallucination and Interpretability

A known challenge with large language models is their tendency to generate plausible but incorrect outputs, often referred to as hallucinations. While our benchmarking focuses on structured biological tasks with ground-truth labels, more open-ended interpretation tasks—such as abstract generation or cluster captioning—may be susceptible to such errors. Developing domain-specific safeguards, such as biological fact-checking mechanisms or constrained decoding strategies, remains an important direction for improving interpretability and reliability in high-stakes settings.

**Figure 8:**
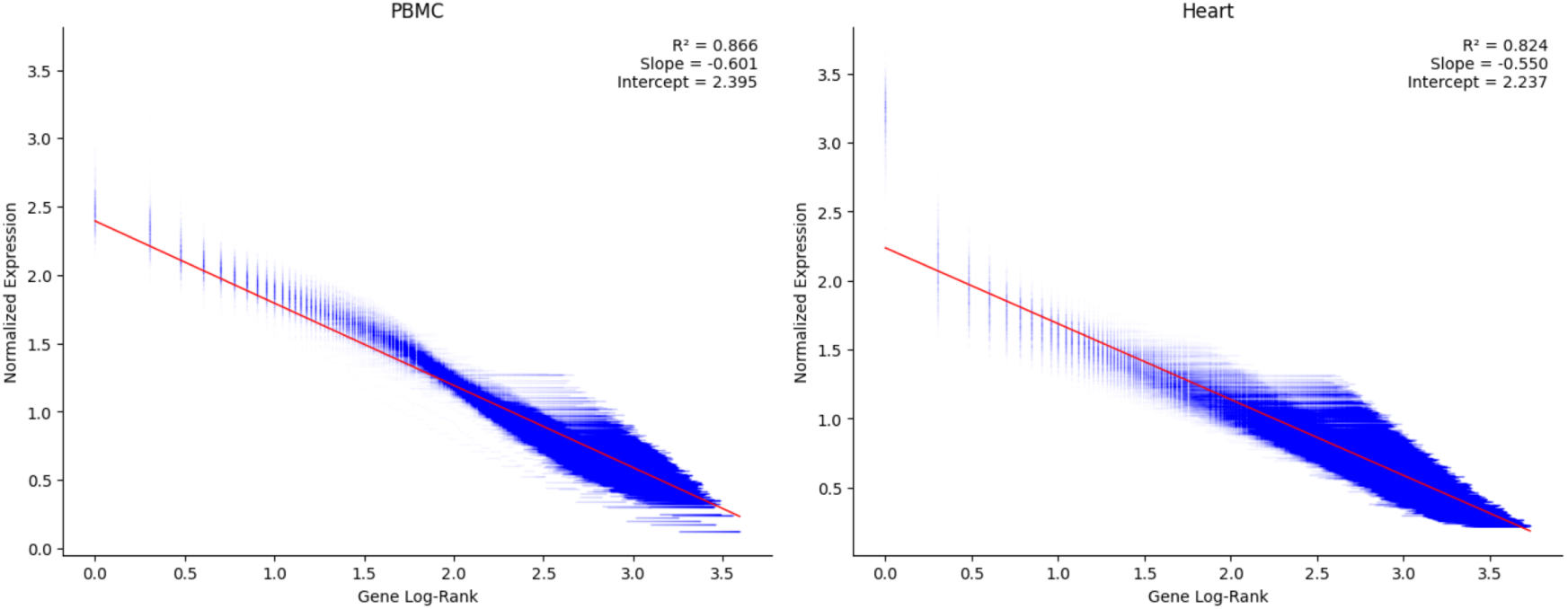
C2S allows for conversion from expression information into cell sentence format with minimal information loss. Using a linear model fitted between rank and original expression, cell sentences can be converted back to expression accurately.

**Figure 9:**
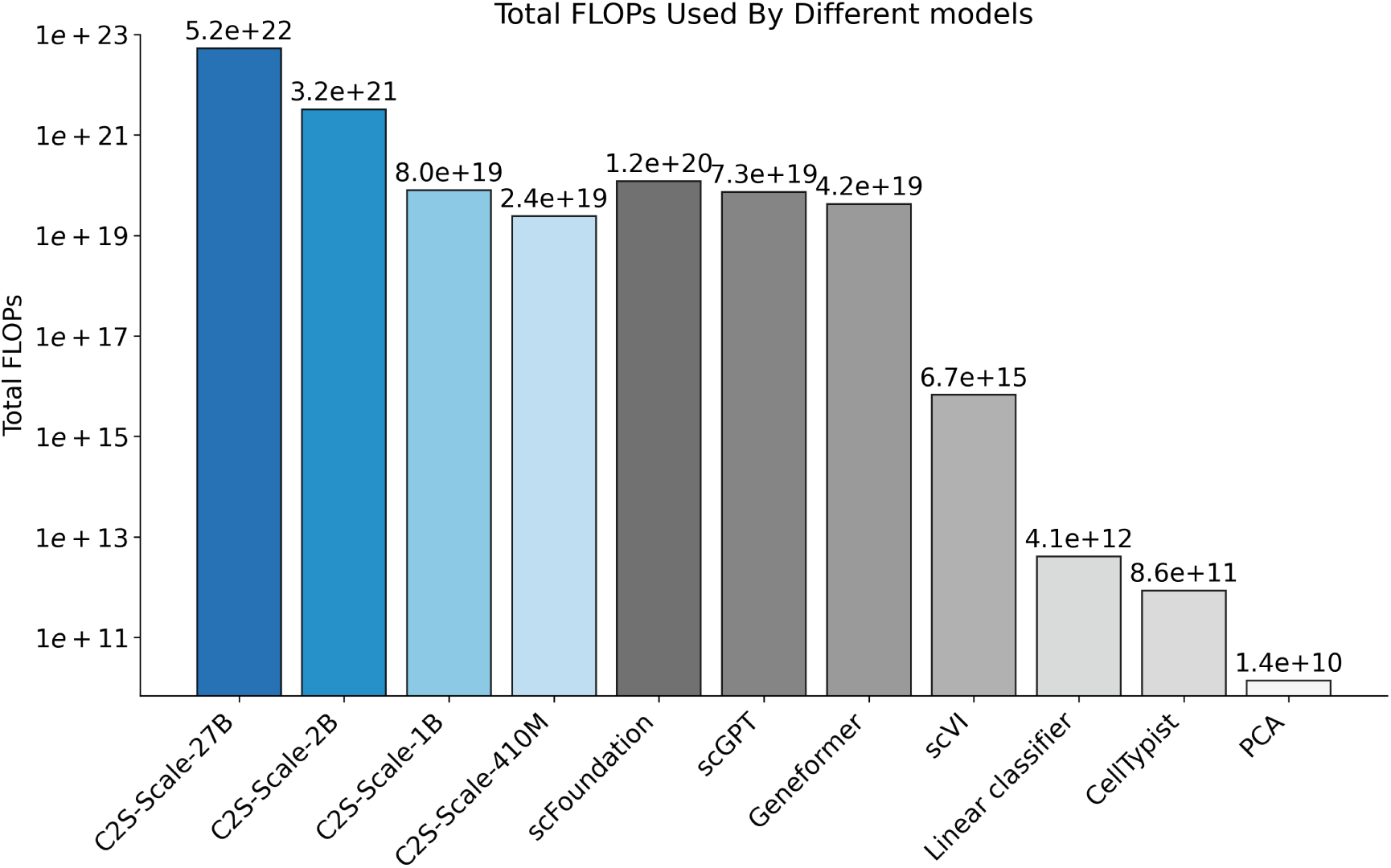
FLOPs usage by C2S-Scale 2B versus other foundation models and simple model baselines.

**Figure 10:**
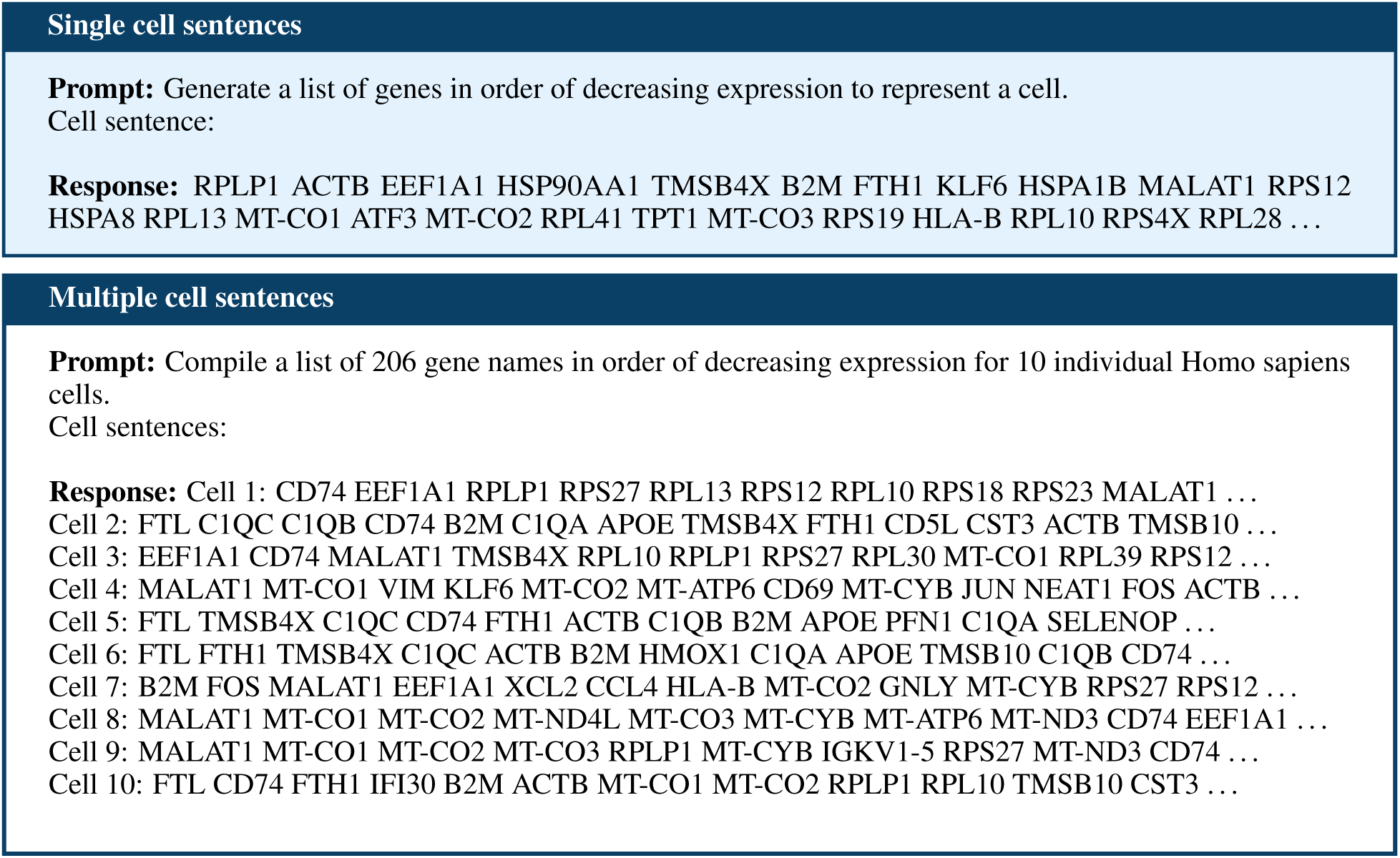
Example input prompts and ground truth responses for C2S-Scale pretraining raw data samples: single-cell and multi-cell samples. Cell sentences are abbreviated for visualization purposes.

**Figure 11:**
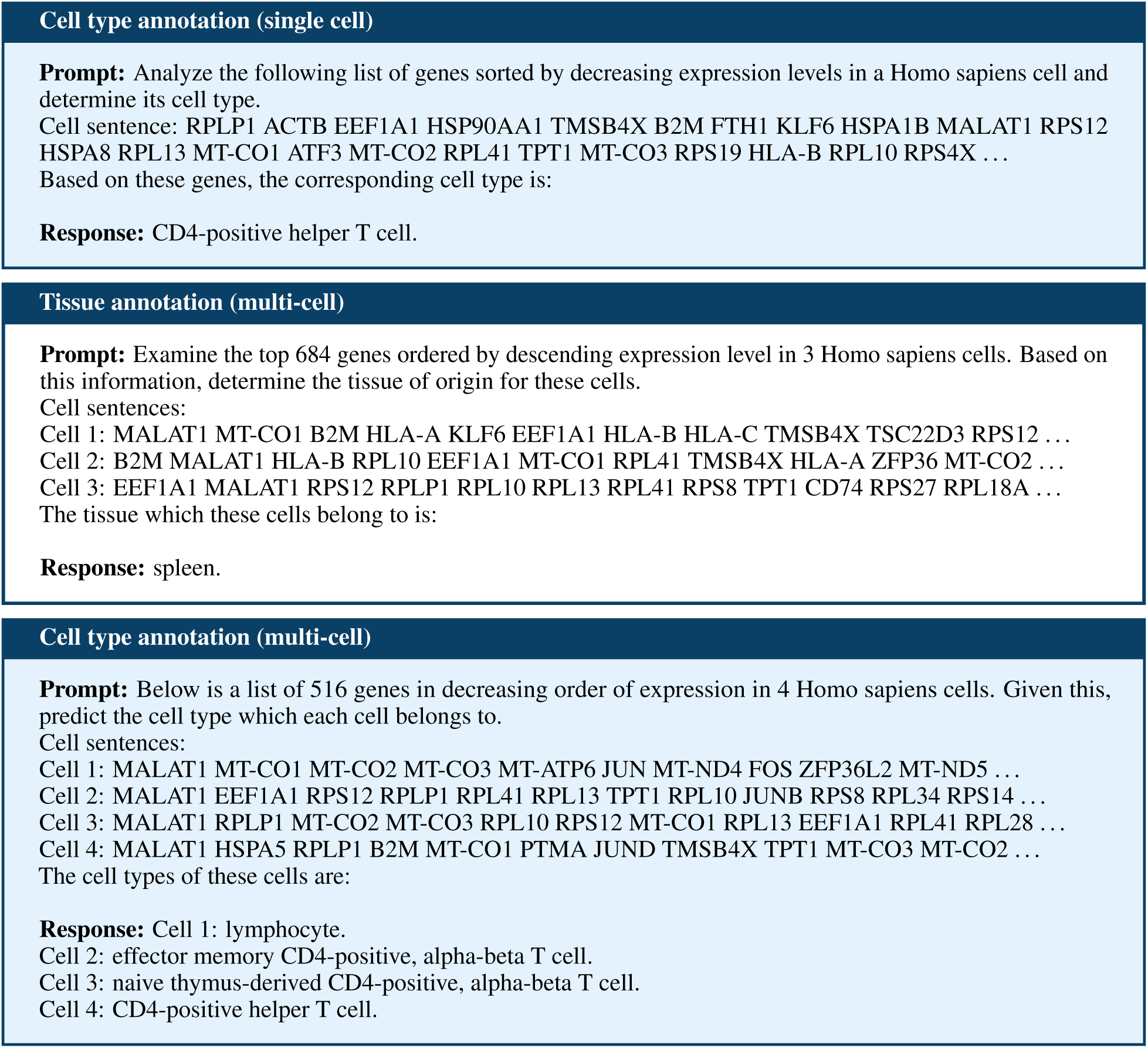
Example input prompts and ground truth responses for C2S-Scale pretraining predictive tasks: cell type(s) annotation and tissue annotation.

**Figure 12:**
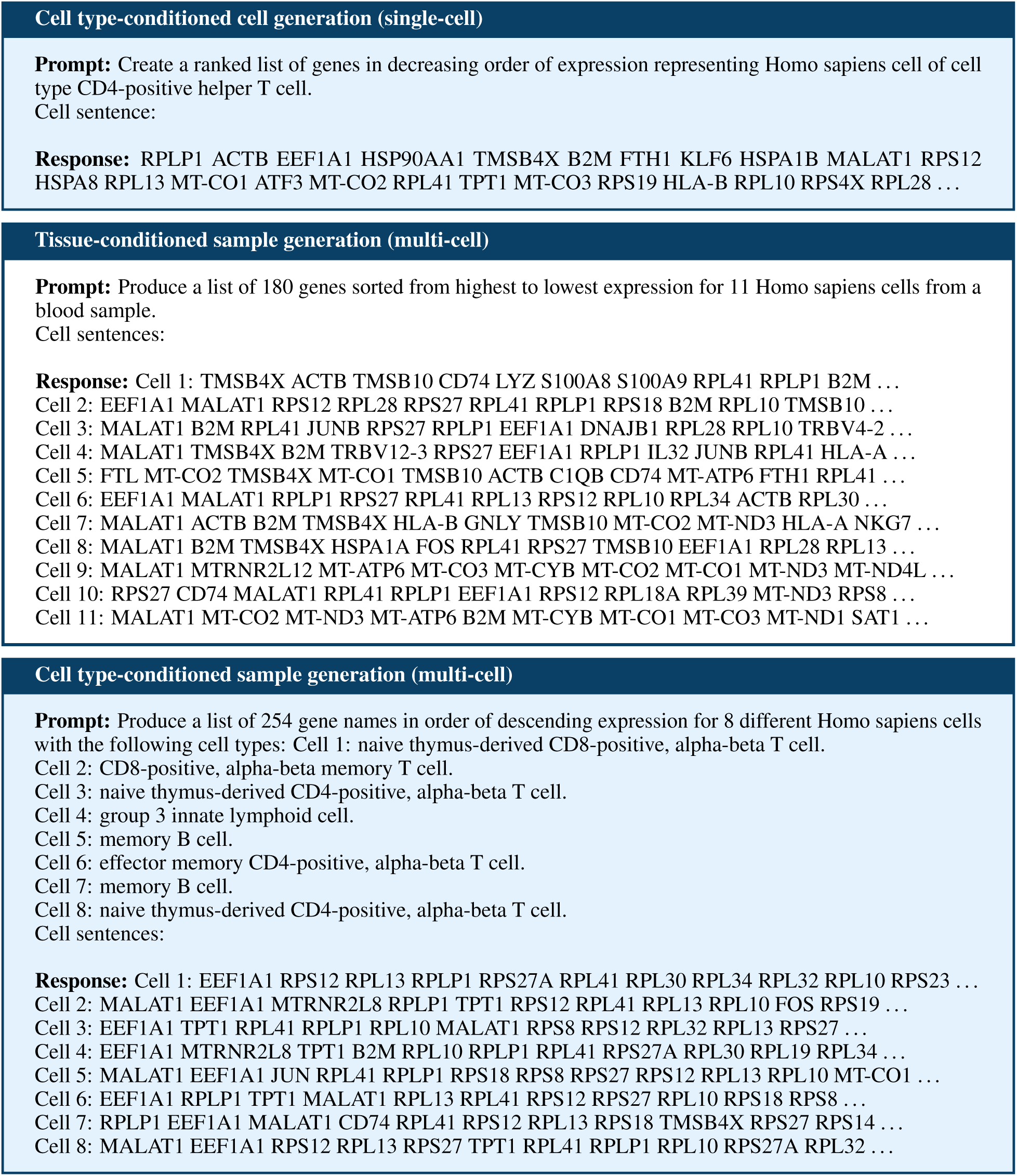
Example input prompts and ground truth responses for C2S-Scale pretraining generation tasks: cell type and tissue conditional generation for single cells and multi-cell samples.

**Figure 13:**
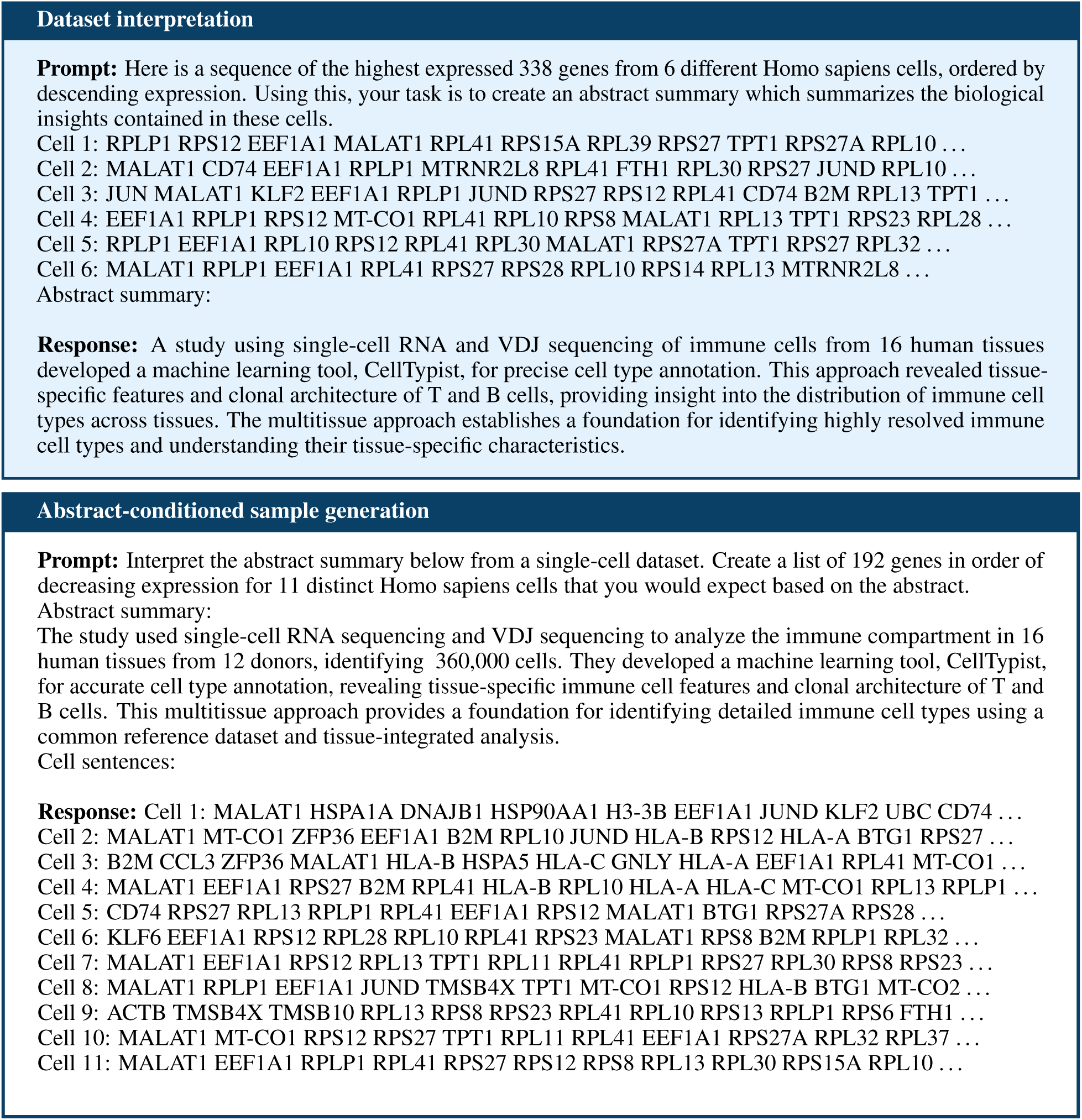
Example input prompts and ground truth responses for C2S-Scale pretraining dataset interpretation tasks: dataset interpretation from multi-cell samples and abstract-conditioned sample generation.

**Figure 14:**
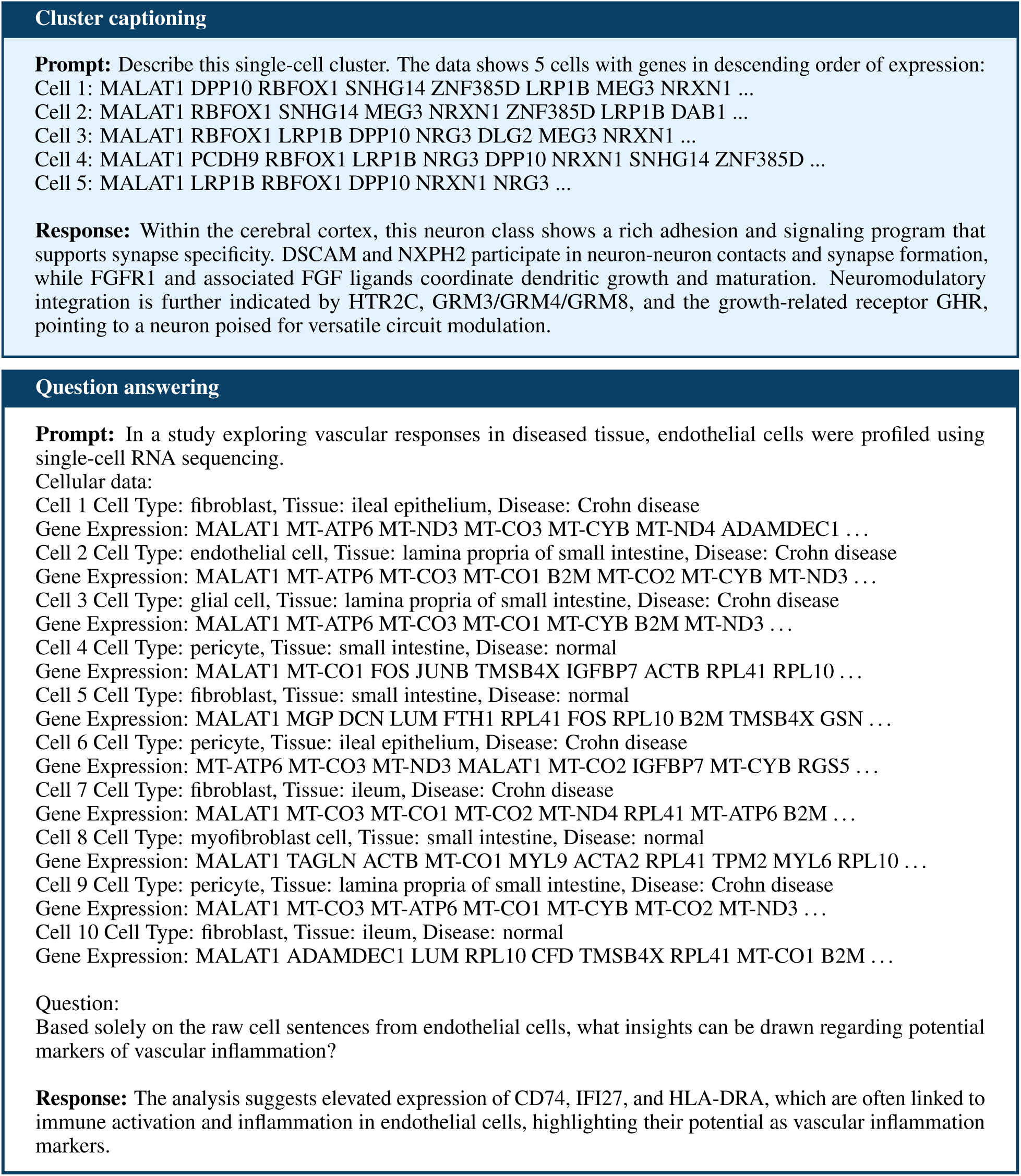
Example input prompts and ground truth responses for C2S-Scale downstream natural language tasks: cluster captioning and question answering.

**Figure 15:**
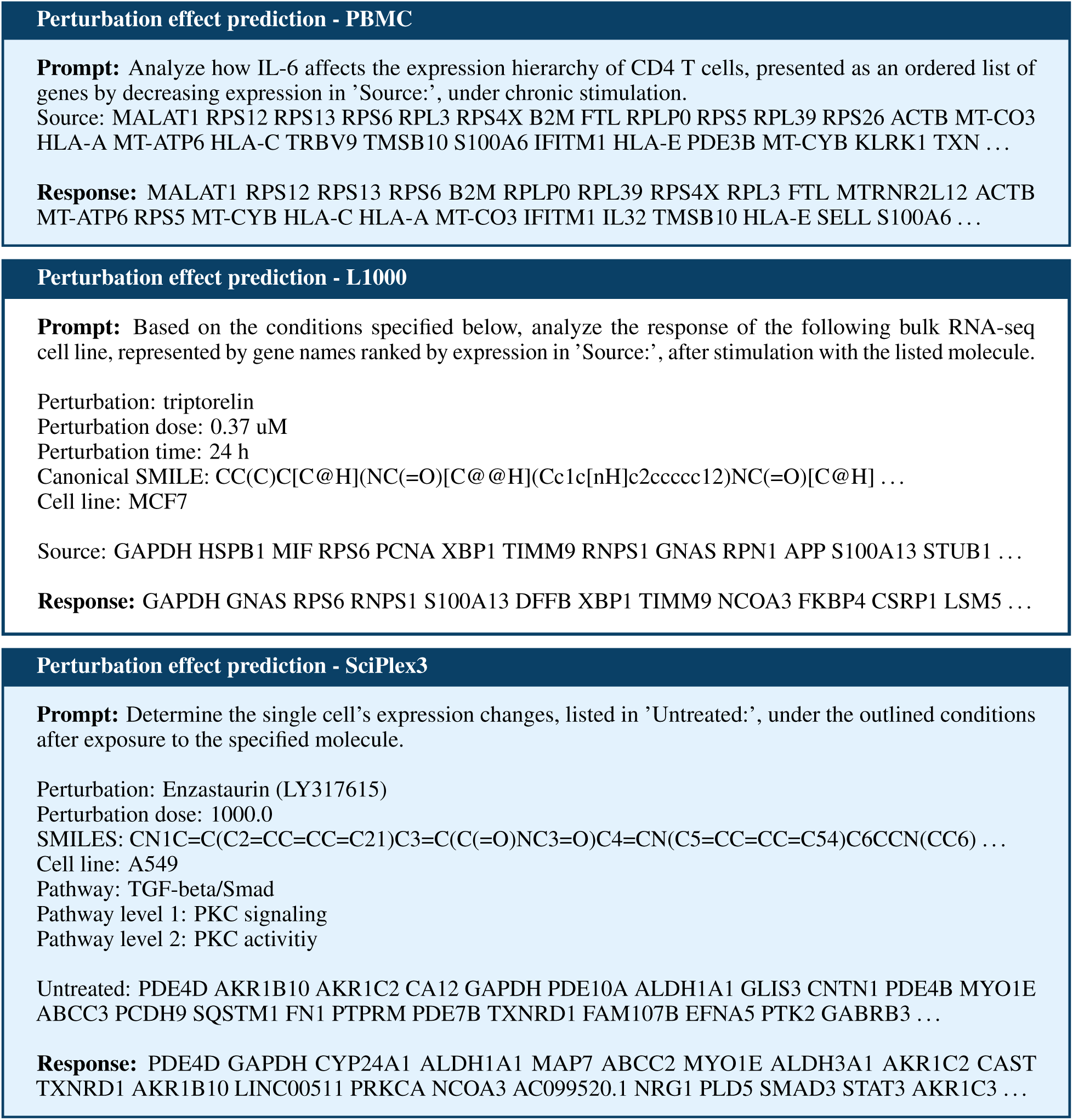
Example input prompts and ground truth responses for C2S-Scale downstream perturbation prediction task on cytokine stimulation PBMC data, L1000, and SciPlex3.

**Figure 16:**
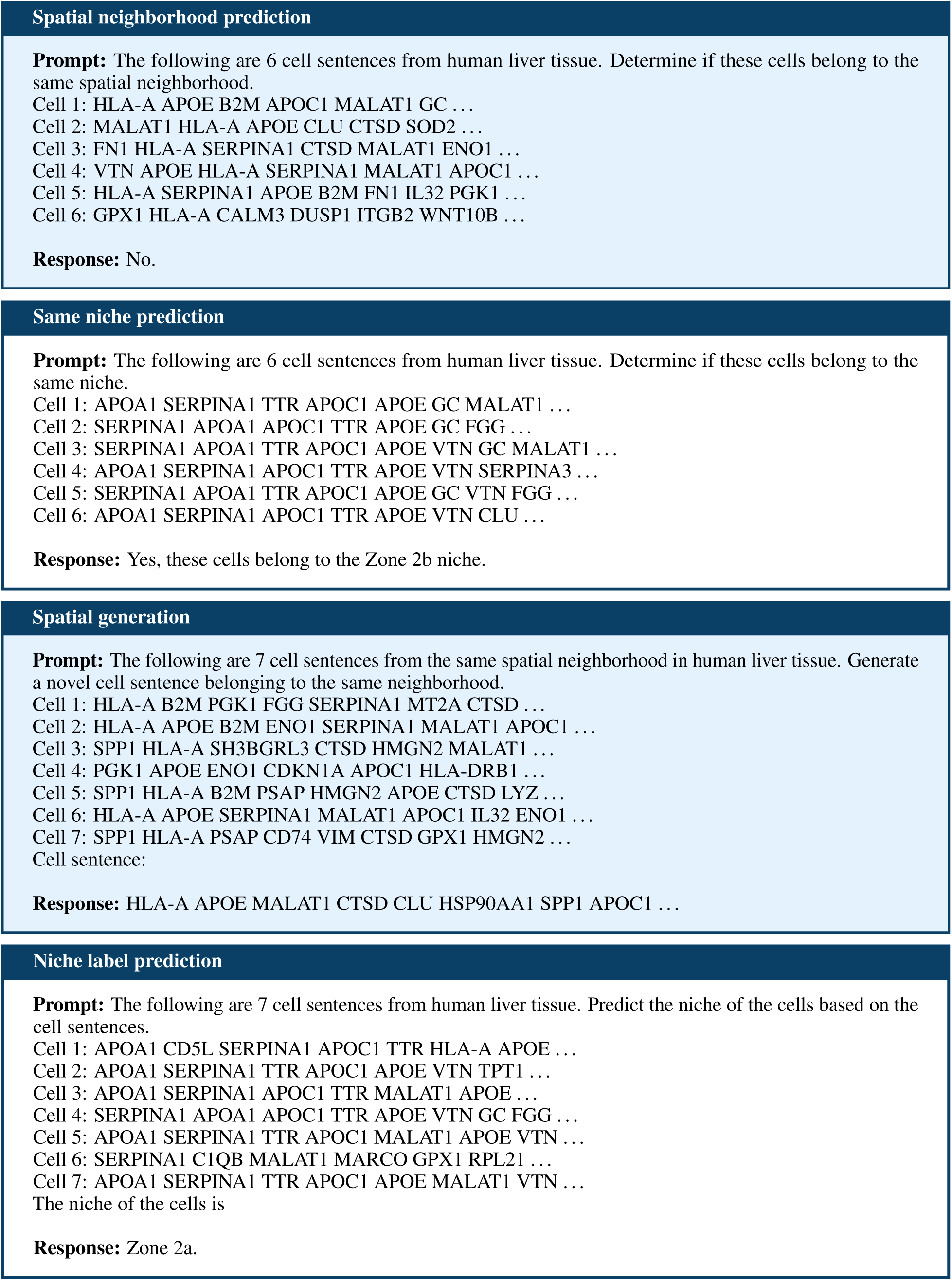
Example input prompts and ground truth responses for C2S-Scale downstream spatial prediction tasks.

**Figure 17:**
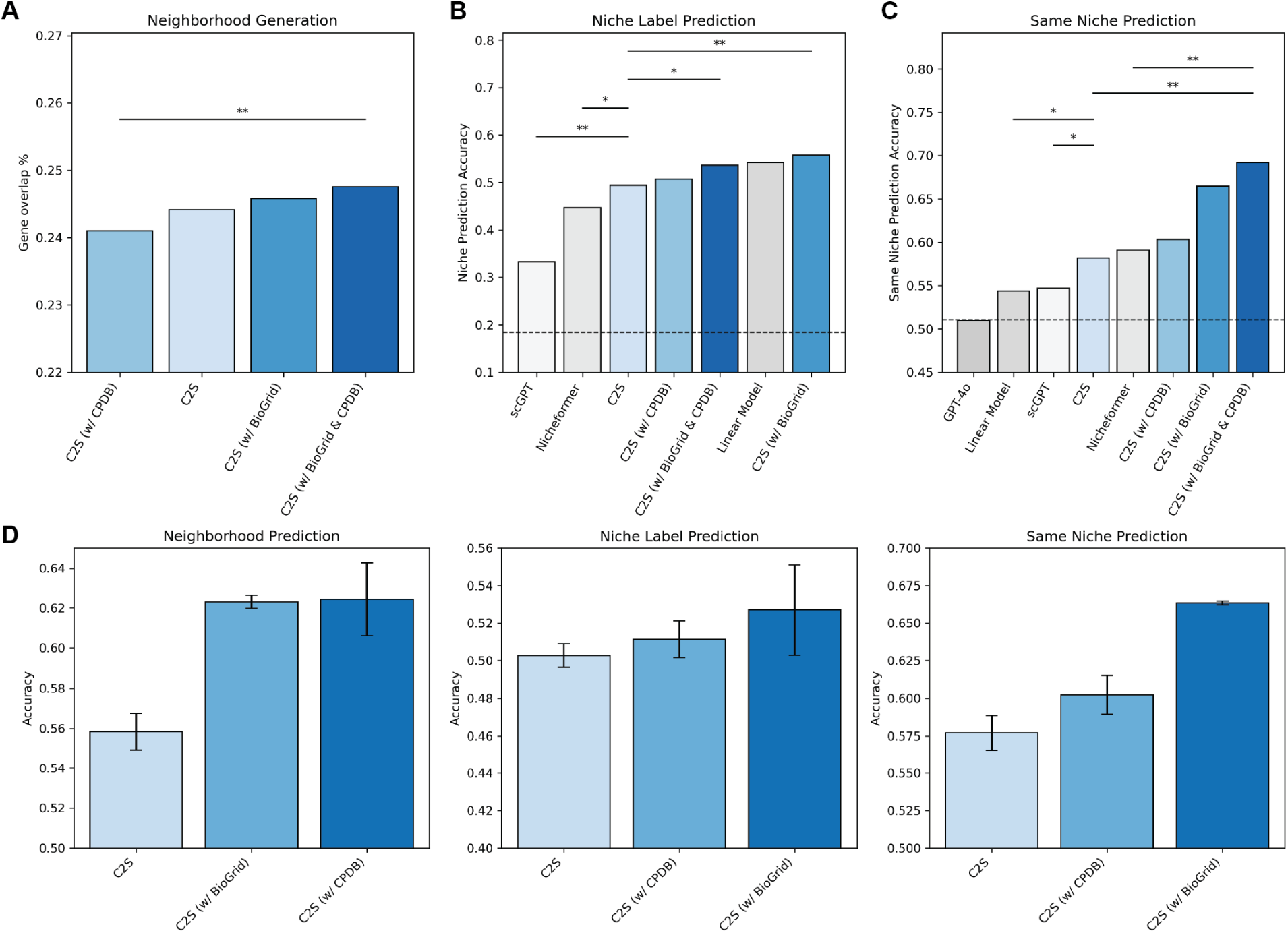
Additional results for spatial tasks. **A,** Spatial cell generation results. Bars indicate average gene overlap percentage across 100 generated replicates for 100 samples. ** = P < 0.01; two-tailed T-test). **B,** Niche label prediction results. Dashed line indicates random accuracy. * = P < 0.05, ** = P < 0.01; McNemar’s test). Dashed line represents random baseline accuracy (0.184). **C,** Same niche prediction results. Dashed line indicates random accuracy. * = P < 0.05, ** = P < 0.01; McNemar’s test). Dashed line represents random baseline accuracy (0.51). **D,** Mean and standard deviation of C2S models trained with three different random seeds. Including CPDB and BioGRID interactions consistently improves performance over the base C2S model.

**Figure 18:**
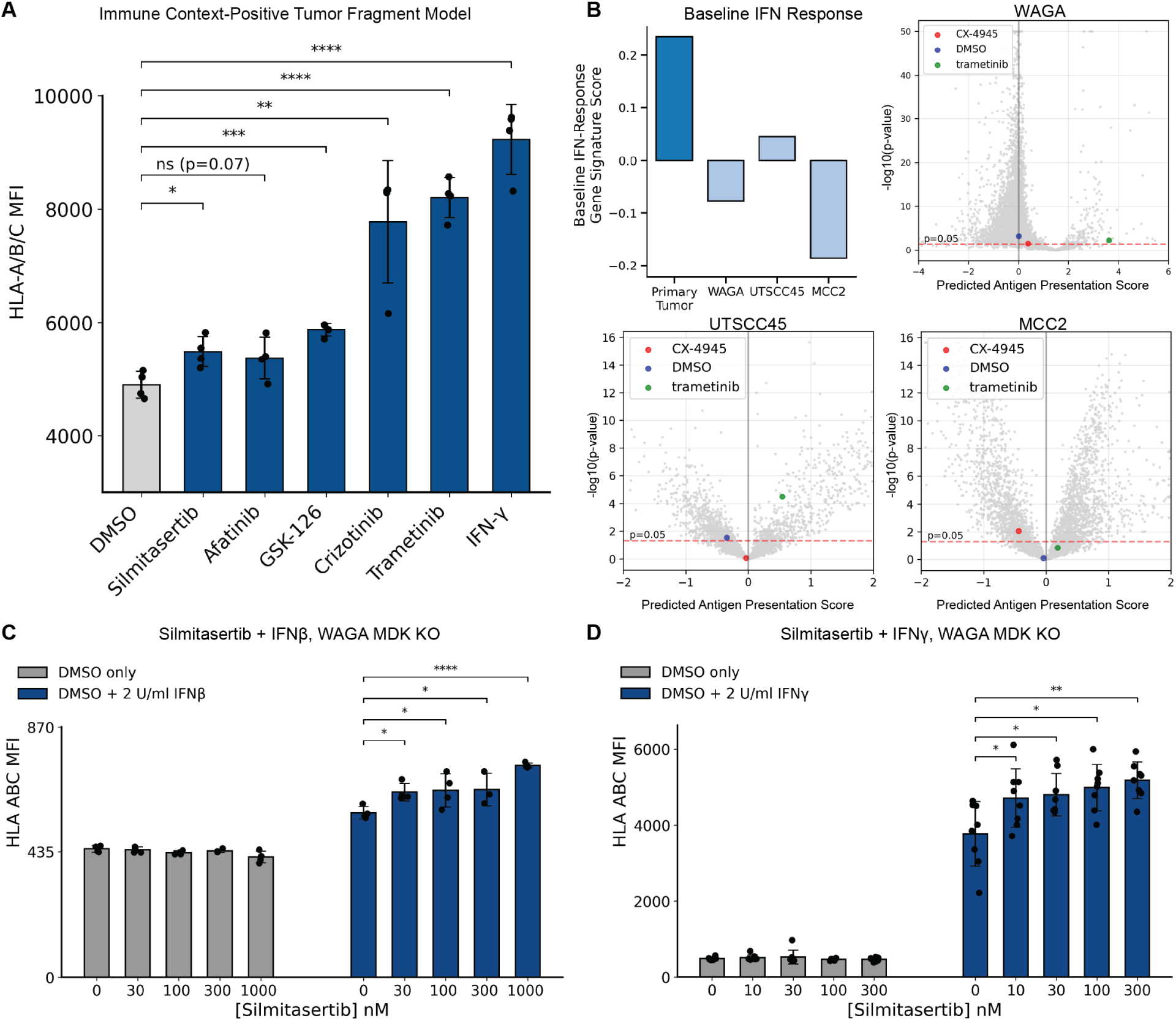
Extended characterization and validation of virtual screening candidates in tumor fragment and cell line models. **A,** Experimental validation of top hits in an immune-context-positive tumor fragment model. Surface levels of HLA-A,B,C (Mean Fluorescence Intensity; MFI) were measured following treatment with 10 nM of each drug. IFN-*γ* serves as a positive control. Significance is compared to DMSO vehicle. **B,** Computational characterization of the immune-context-neutral state. (Top Left) Baseline Type I interferon (IFN) response gene signature scores for the Primary Tumor context compared to three cell lines (WAGA, UTSCC45, MCC2). (Top Right, Bottom) Volcano plots displaying predicted drug effects in these cell line contexts. Silmitasertib (red marker) shows no significant predicted efficacy in these interferon-low settings compared to the primary tumor context. **C,** Validation of the interferon-conditional effect in MDK-knockout WAGA cells. Dose-response assessment of HLA-A,B,C MFI in MDK-knockout WAGA cells treated with silmitasertib in the absence (gray) or presence (blue) of low-dose IFN-*β* (2 U/ml). **D,** Generalization of the amplification effect to Type II interferon. Dose-response assessment of HLA-A,B,C MFI in MDK-knockout WAGA cells treated with silmitasertib in the absence (gray) or presence (blue) of low-dose IFN-*γ* (2 U/ml). Error bars represent mean ± s.d. Significance calculated via two-way t-test with Benjamini-Hochberg correction; ∗*P <* 0.05, ∗ ∗ *P <* 0.01, ∗ ∗ ∗*P <* 0.001, ∗ ∗ ∗ ∗ *P <* 0.0001.

**Figure 19:**
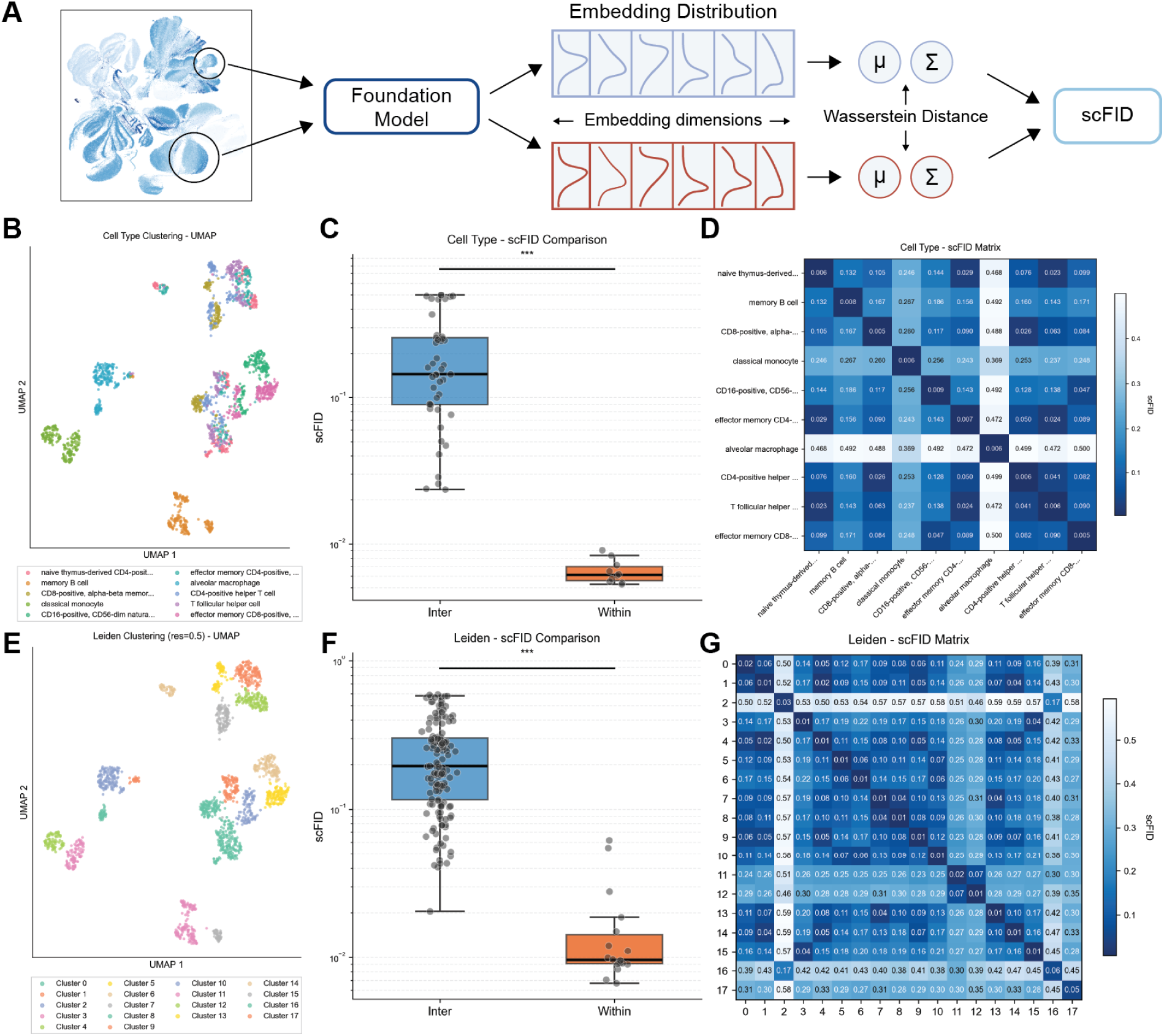
**A,** Diagram of the scFID metric, computed in foundation model latent space, analogous to FID in computer vision. **B,** UMAP showing the 10 most frequent cell types in the Dominguez immune dataset [20]. **C,** The scFID is significantly lower within cells of the same cell type than between different cell types (two sample t-test; 10 samples within cell types, 45 samples between cell types; *P* = 2.23 × 10^−4^). **D,** Confusion matrix showing the scFID between every pair of cell types. **E-G** Same as B-D, but for Leiden clusters instead of cell (two sample t-test; 18 samples within clusters, 153 samples between clusters; *P* = 2.60 × 10^−9^.)

**Figure 20:**
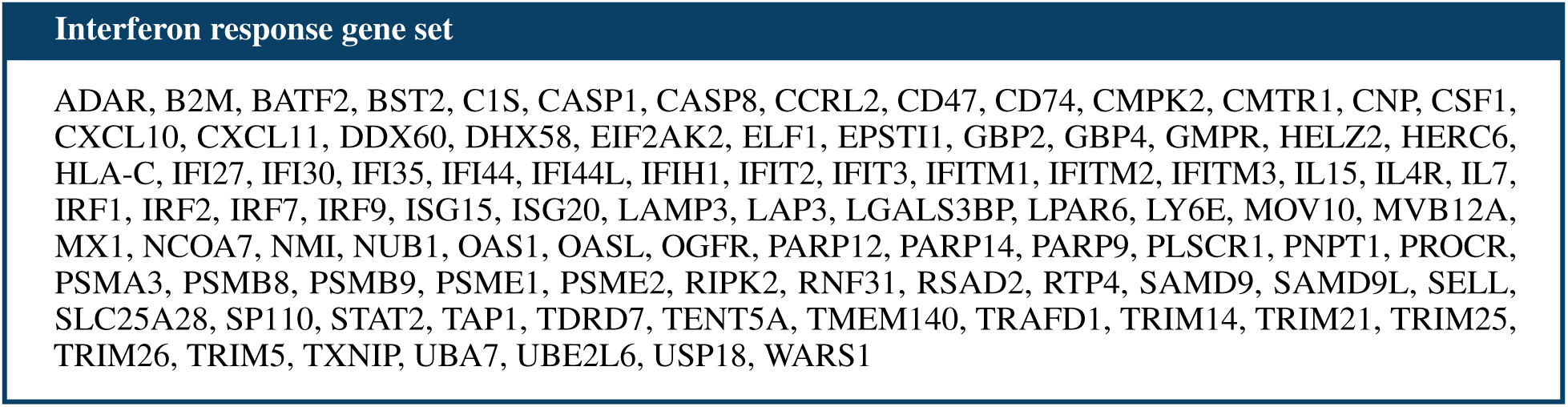
The 97 genes in the “Hallmark Interferon Alpha Response” gene set, used as a proxy for estimating the level of baseline interferon activity for virtual screening.

**Figure 21:**
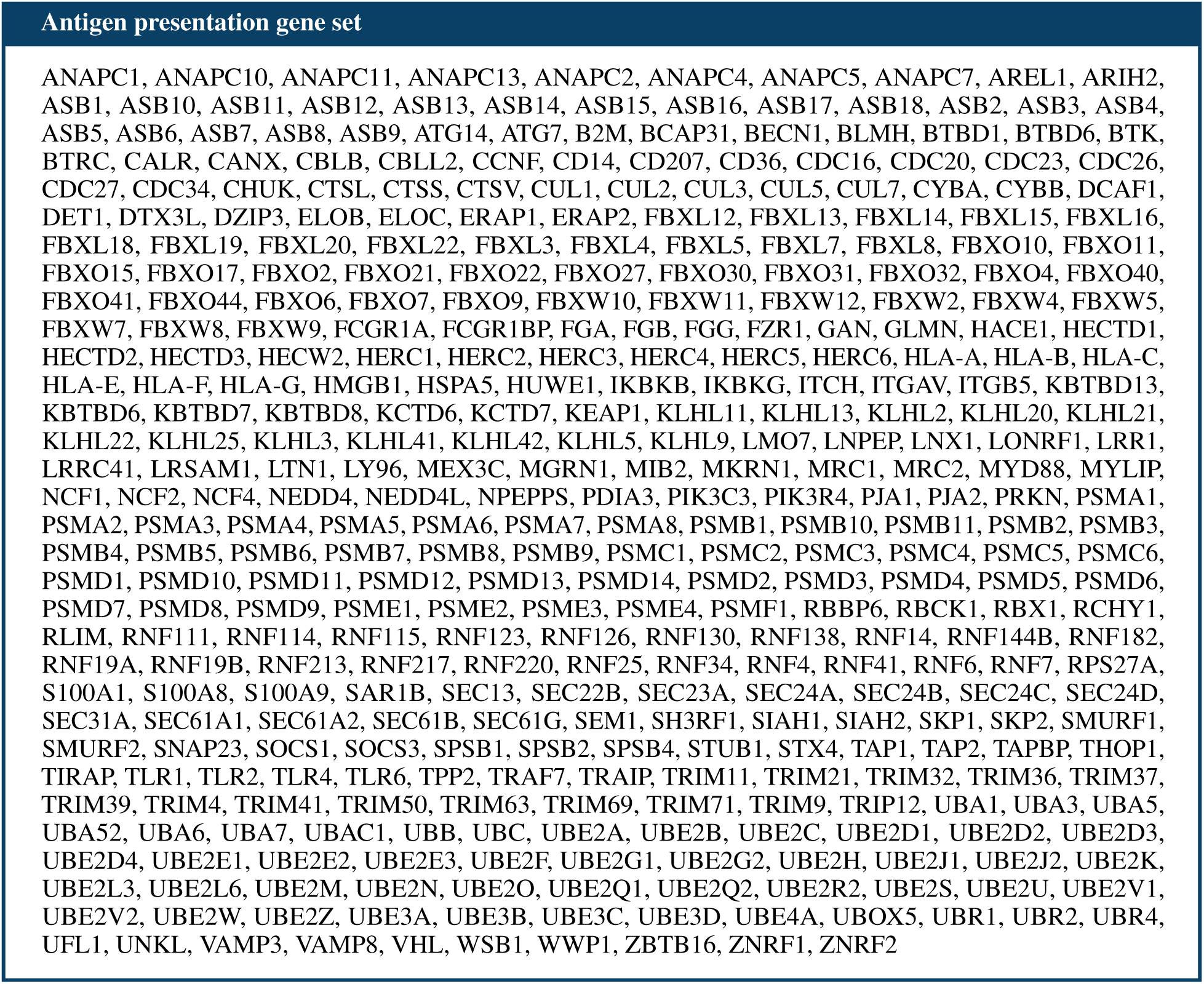
The 381 genes in the “Reactome Class I MHC-Mediated Antigen Processing Presentation” gene set, used to score virtual screen outputs.

